# PCBP1 regulates alternative splicing of AARS2 in congenital cardiomyopathy

**DOI:** 10.1101/2023.05.18.540420

**Authors:** Yao Wei Lu, Zhuomin Liang, Haipeng Guo, Tiago Fernandes, Ramon A Espinoza-Lewis, Tingting Wang, Kathryn Li, Xue Li, Gurinder Bir Singh, Yi Wang, Douglas Cowan, John D Mably, Caroline C. Philpott, Hong Chen, Da-Zhi Wang

## Abstract

Alanyl-transfer RNA synthetase 2 (AARS2) is a nuclear encoded mitochondrial tRNA synthetase that is responsible for charging of tRNA-Ala with alanine during mitochondrial translation. Homozygous or compound heterozygous mutations in the Aars2 gene, including those affecting its splicing, are linked to infantile cardiomyopathy in humans. However, how Aars2 regulates heart development, and the underlying molecular mechanism of heart disease remains unknown. Here, we found that poly(rC) binding protein 1 (PCBP1) interacts with the Aars2 transcript to mediate its alternative splicing and is critical for the expression and function of Aars2. Cardiomyocyte-specific deletion of Pcbp1 in mice resulted in defects in heart development that are reminiscent of human congenital cardiac defects, including noncompaction cardiomyopathy and a disruption of the cardiomyocyte maturation trajectory. Loss of Pcbp1 led to an aberrant alternative splicing and a premature termination of Aars2 in cardiomyocytes. Additionally, Aars2 mutant mice with exon-16 skipping recapitulated heart developmental defects observed in Pcbp1 mutant mice. Mechanistically, we found dysregulated gene and protein expression of the oxidative phosphorylation pathway in both Pcbp1 and Aars2 mutant hearts; these date provide further evidence that the infantile hypertrophic cardiomyopathy associated with the disorder oxidative phosphorylation defect type 8 (COXPD8) is mediated by Aars2. Our study therefore identifies Pcbp1 and Aars2 as critical regulators of heart development and provides important molecular insights into the role of disruptions in metabolism on congenital heart defects.

## INTRODUCTION

Congenital heart disease (CHD) is the most common category of birth defects and a leading cause of morbidity and mortality in children worldwide. While the genetic causes of certain types of CHD have been identified, the molecular basis for the rest remains elusive. RNA-binding proteins (RBPs) play critical roles during cardiac development^1,2^, and are important post-transcriptional regulators for RNA metabolism involved in alternative-splicing, RNA stability, and translation processes. While RBPs have been implicated in the development and maturation of the heart^1,3,4^, the molecular and biological roles for many RBPs in heart development have not been determined.

Following post-transcriptional modification, a crucial step in protein synthesis is the ‘charging’, or aminoacylation of transfer RNA (tRNA), by tRNA synthetase. There are cytosolic and mitochondrial tRNA synthetases that largely function in separate cellular compartments, but all are encoded in the nucleus. Intriguingly, all mitochondrial tRNA synthetases are associated with autosomal recessive human disorders that manifest in tissue- and cell-specific phenotypes, despite their ubiquitous expression patterns^5^. The recessively inherited alanyl-tRNA synthetase 2 (AARS2) mutation was first described in patients presenting with infantile cardiomyopathy with defects in oxidative phosphorylation^6^. Later on, the spectrum of AARS2-related disease expanded to include leukodystrophy spectrum disorders. The different spectra of AARS2-related disease has been attributed to the different locations of pathogenic variants in the protein and their effects on protein function^7^. The importance of Aars2 is further supported by knock-in mice harboring editing domain mutations, resulting in early embryonic lethality prior to E8.5 in homozygous mutants^8^. Due to the early lethality in mice with mutations in the AARS2 editing domain, it remains challenging to study Aars2 in the context of heart development *in vivo*.

PCBP1 belongs to the poly(rC)-binding proteins (PCBP) family of RBPs; there is only one member in invertebrates but four members in mammalians (Pcbp1, Pcbp2, Pcbp3, and Pcbp4)^9,10^. The intron-less Pcbp1 gene is highly conserved in mammals and is thought to have evolved from Pcbp2 through a retrotransposition event, deduced based on the stringent structural conservation of the Pcbp1 locus; however, it has critical but distinct cellular functions^10,11^. The PCBP family of RBPs plays important roles in posttranscriptional RNA processing, alternative-splicing, and translation regulation by interacting with pyrimidine-rich motifs in targeted RNAs^9,10, 12–14^, and also serves unconventional functions such as transcription^12^ and iron trafficking through ferritin^15–17^. Perturbation of PCBP paralogs in invertebrates such as Drosophila^18^ and C. elegans^19^ leads to lethal phenotypes. Germline deletion of either Pcbp1 or Pcbp2 in mice is embryonically lethal^20,21^, suggesting that they are independently essential during development. Conditional inactivation of Pcbp1 or Pcbp2 individually in the erythroid lineage using EpoR-Cre failed to significantly impact the development of the erythroid lineage, while the combined conditional inactivation of Pcbp1 and Pcbp2 results in the loss of hematopoiesis and blunted erythroid development. These results suggest that Pcbp1 and Pcbp2 serve redundant roles in the development of the erythroid lineage^22^. Interestingly, despite the redundancy of Pcbp1 in erythroid development, the expression of Pcbp1 alone is sufficient for erythroid heme and globin production postnatally^17^. Mice with inducible depletion of Pcbp1 in adulthood with Rosa26-CAG-CreERT2 exhibited microcytic anemia and activation of compensatory erythropoiesis^17^. Pcbp1 is also critical for effector T-cell differentiation and serves as an intracellular immune checkpoint for T-cell function and cancer immunity^23^. These studies underscore the critical function that Pcbp1 governed in these cell-types independent of Pcbp2. We have previously generated a hypomorphic Pcbp1-mutant mouse line that displays defects in skeletal muscle growth due to defects in the proliferation and differentiation of skeletal muscle myoblasts and satellite cells, through modulation of the microRNA processing pathway^21^. Pcbp1 had been identified as one of the heterogeneous nuclear ribonucleoproteins that regulates collagen synthesis at the post-transcriptional level by interacting with the 3’-UTR of the mRNA in the cardiac fibroblast^24^. However, the function of Pcbp1 in cardiomyocytes, especially during cardiac development, is largely unexplored.

In this study, we sought to determine the mechanistic role of Pcbp1 and Aars2 during heart development. Using genetic and biochemical approaches, we discovered an essential role of Pcbp1 in heart development through regulation of alternative splicing of Aars2, which is essential for Aars2 function. Genetic deletion of either Pcbp1 or Aars2 in cardiomyocytes leads to noncompaction cardiomyopathy, as well as a consistent and significant reduction of proteins that constitute the oxidative phosphorylation complexes. Our findings uncovered a novel function of Pcbp1 in regulating Aars2 alternative splicing in cardiomyocytes during development, which may offer new insights into the RBP-regulated mechanisms of CHD.

## RESULTS

### Pcbp1 interacts with Aars2 mRNA near the splice site of exon 16 in the heart

Pathogenic mutations of AARS2 located in the editing domain have been identified as causal for lethal infantile cardiomyopathy^6,7,25^. Additional damaging variants located within the AARS2 editing domain have been recently identified through whole genome and exon studies^26^, including two intronic pathogenic variants located at the splice donor site (c2255+1G>A) and splice acceptor site (c2146-2A>G) flanking exon 16 of AARS2 (**Supplemental Figure 1A**). These mutations disrupt the canonical donor (GT) or acceptor (AG) sites and are predicted to result in the skipping of exon 16. The individual with these variants has clinical features of COXPD8, including cardiomyopathy (**Supplemental Figure 1B**). This evidence suggests a potential link of the AARS2-related infantile cardiomyopathy clinical feature and the alternative splicing of AARS2 at exon 16.

We hypothesized that RBPs interacting with the AARS2 mRNA near exon 16 may play a role in directing its proper alternative splicing. We performed an *in silico* screen using RBPsuite^27^ with the pre-mRNA sequence from exon 15 to exon 17 of AARS2 to look for enrichment of consensus RBP motifs, and we found that the consensus PCBP1 motifs are highly enriched at the intron-exon junction of exon 16, as well as within exons 15 and 17 (**Figure 1A**).

**Figure 1.**
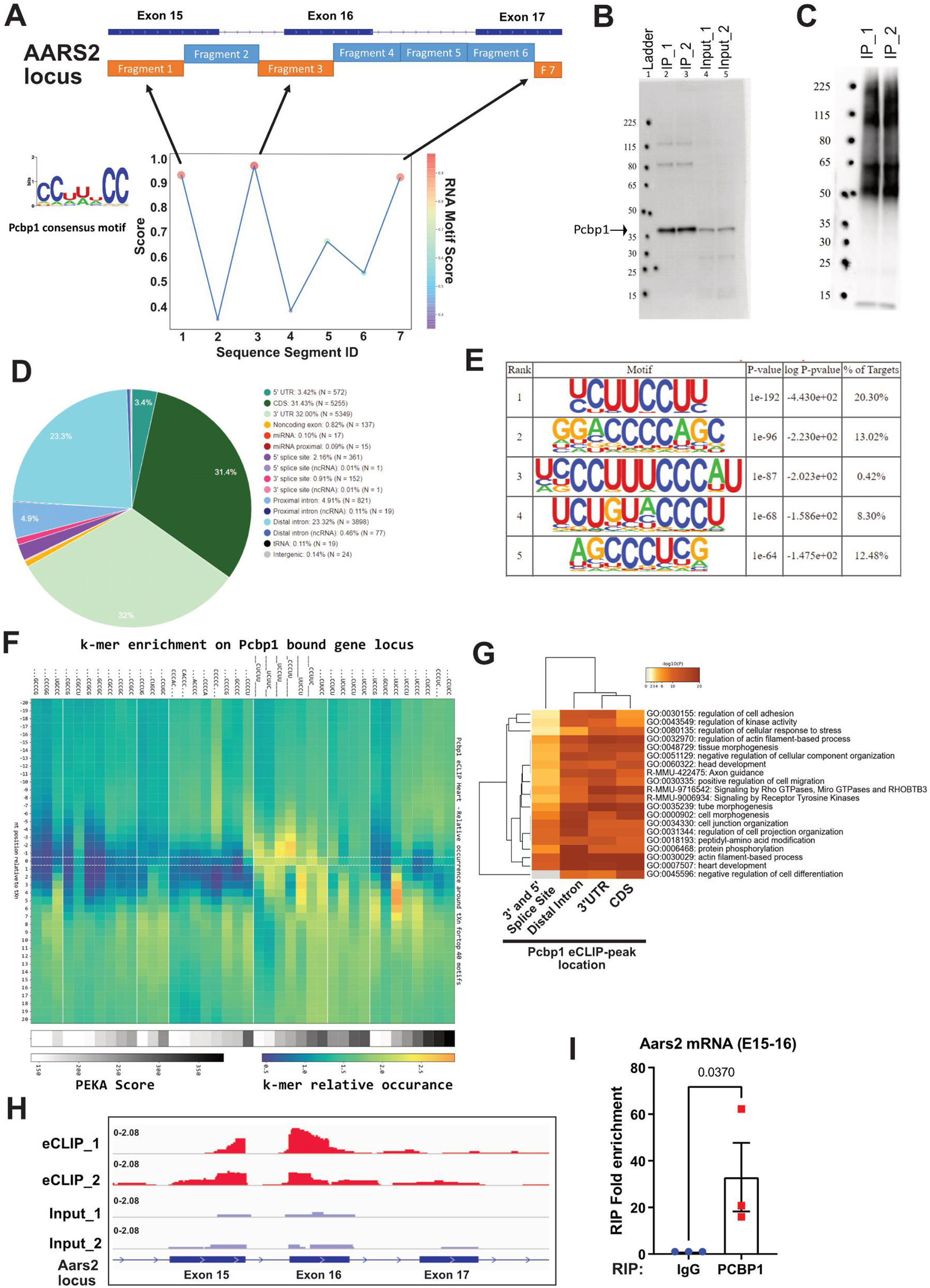
PCBP1 interacts with Aars2 mRNA transcript in the heart during development. (A) In silico motif analysis revealed an enrichment of Pcbp1 consensus motif near exon 16. (B) Enhanced cross-linked immunoprecipitation followed by sequencing (eCLIP-seq) of PCBP1 were performed with E16.5 embryonic heart ventricles (18-19 hearts per replicate for two replicates). Western blotting of PCBP1 were performed after eCLIP, only regions from 40 kDa to 115 kDa were isolated during eCLIP. (C) PCBP1-bound RNA was visualized on nitrocellulose membrane during eCLIP in embryonic heart tissue. (D) eCLIP-seq bound peak regions were further classified, and the relative frequency of peaks that map to each feature type were depicted in a pie chart, with a peak log2 fold enrichment ≥ 3 and p-value ≤ 0.001. (E) Enriched motifs in eCLIP-seq peaks were identified by de novo motif discovery algorithm HOMER, and the top 5 enriched motifs were shown. (F) Positionally enriched k-mer analysis (PEKA) was performed, and the k-mer enrichment relative to the cross-linked site of the eCLIP shown. (G) Gene ontology (GO) enrichment for the genes with PCBP1 interactions at the 3’ and 5’ splice sites, distal intron, 3’UTR, and CDS were shown. (H) eCLIP-seq peaks for PCBP1 located near the exon 15 to exon 17 region is shown. (I) RNA immunoprecipitation (RIP) assay was performed with PCBP1 antibody on neonatal heart ventricle lysates, and Aars2 primers spanning the exon 15 to exon 16 regions were assayed by qRT-PCR on RIP-bound RNA (n=3, Student’s T-Test p-value shown,, data presented as mean ± SEM).

To assess and identify the direct RNA-binding of Pcbp1 in the developing heart, we performed enhanced UV crosslinking and immunoprecipitation followed by high-throughput sequencing (eCLIP-seq)^28^ with wild-type E16.5 hearts. We confirmed successful eCLIP immunoprecipitation of PCBP1 (**Figure 1B**) by RNA immunoprecipitation (RIP) (**Figure 1C**). Irreproducible discovery rate (IDR) PCBP1 peaks were identified using the previously described pipeline CLIPper ^28,29^ and is consistent with the Encyclopedia of DNA Element (ENCODE) standards. In our PCBP1 eCLIP-seq data, we mapped the location of the interaction in the context of the transcriptome and found that 32% of the PCBP1 binding was located at the 3’UTR, while 31.43% and 23.32% were located in the coding sequence (CDS) and distal intron regions, respectively (**Figure 1D**). The 3’UTR bias in its binding is apparent and consistent with a previous report in human cells^30^, but the relatively high enrichment of binding in the CDS and distal intron regions suggests its propensity for regulating alternative splicing and transcript stability^31^.

The RNA motifs identified to be bound by PCBP1 in the heart are pyrimidine-rich with stretches of C and U repeats (**Figure 1E**), consistent with PCBP1 binding motif-patterns identified in cell-free systems and in vitro human cell lines ^30,32^. Through positional motif k-mer analysis^33^, we found that the pyrimidine-rich k-mer was preferentially enriched at least 5-nucleotides downstream of the eCLIP cross-link site (**Figure 1F, Supplemental Figure 4A-C**). Focusing on the PCBP1 binding peaks identified at the binding regions (i.e. 3’ and 5’ splice sites, distal intron, 3’UTR, and CDS), we found the transcript loci of these binding regions to be co-enriched for GO-terms of developmental processes such as “heart development”, “cell morphogenesis”, “tissue morphogenesis,” and “regulation of cellular response to stress” (**Figure 1G**). Interestingly, 40-50% of these transcript loci contained multiple PCBP1 binding peaks in different transcript regions, revealing a more complex and potentially combinatorial regulation of these transcripts by Pcbp1. Importantly, we identified significant interaction of PCBP1 with AARS2 mRNA at the exon 15 to exon 16 region (**Figure 1H and I**), further suggesting the potential function for PCBP1 in regulating AARS2.

### Genetic deletion of Pcbp1 in cardiomyocytes led to noncompaction cardiomyopathy and bifurcated apex formation

Pcbp1 was highly expressed in the embryonic heart, and gradually decreased in expression in the adult heart (**Figure 4A**). To study the function of Pcbp1 in the developing heart, we generated myocardial-specific deletion of Pcbp1 (Pcbp1-cKO) by crossing Pcbp1-Flox mice^17,21^ with Ctnt-Cre^34^. Pcbp1-cKO appeared in expected Mendelian ratios at embryonic stage (E) 12.5, E16.5, and postnatal day (P) 1. However, Pcbp1-cKO at weaning age of P28 were underrepresented, suggesting perinatal lethality (**Supplemental Figure 2A-E**). During development prior to extensive coronary formation, the myocardial wall undergoes remodeling characterized by the formation of cardiac trabeculae as a means to increase nutrient exchange surface area and increase cardiomyocyte mass, to meet the increasing metabolic demands of the heart. Between E12.5 and E16.5, as the coronary plexus begins to cover the ventricles to support the myocardial wall, the base of the cardiac trabeculae begins to thicken and become indistinguishable from the myocardium^35^. As the compact zone of the myocardium grows, it replaces the trabeculae as the major contractile force; the failure of this often results in left ventricular non-compaction (LVNC), a structural and functional defect of the heart^36^. To investigate the morphological consequences of the myocardial deletion of Pcbp1, we examined the embryonic heart with brightfield dissecting microscopy followed by H&E staining at E12.5 (**Figure 2A,D**), E16.5 (**Figure 2B,E**), and P1 (**Figure 2C,F**). The Pcbp1-cKO heart displayed abnormal ventricular development signified by the bifurcated apex at E16.5 (**Figure2 B,E**). Additionally, the ventricular compact zone of Pcbp1-cKO heart maintained a reduced thickness in both E16.5 and P1. We then examined the embryonic heart at E12.5 by high-resolution episcopic microscopy of the whole embryo, followed by 3D reconstruction to assess the ventricular morphology (**Figure 2G,H**)^37^, and found that the Pcbp1-cKO heart displayed shorter statute with higher width to height ratio (**Figure 2I-K**) and markedly thinner compact zone in the left and right ventricles (**Figure 2L-M**). Together with the hypertrabaculation and hypoplasia, these are hallmarks indicative of ventricular non-compaction defects, which persisted at P1 (**Figure 2C,F**).

**Figure 2.**
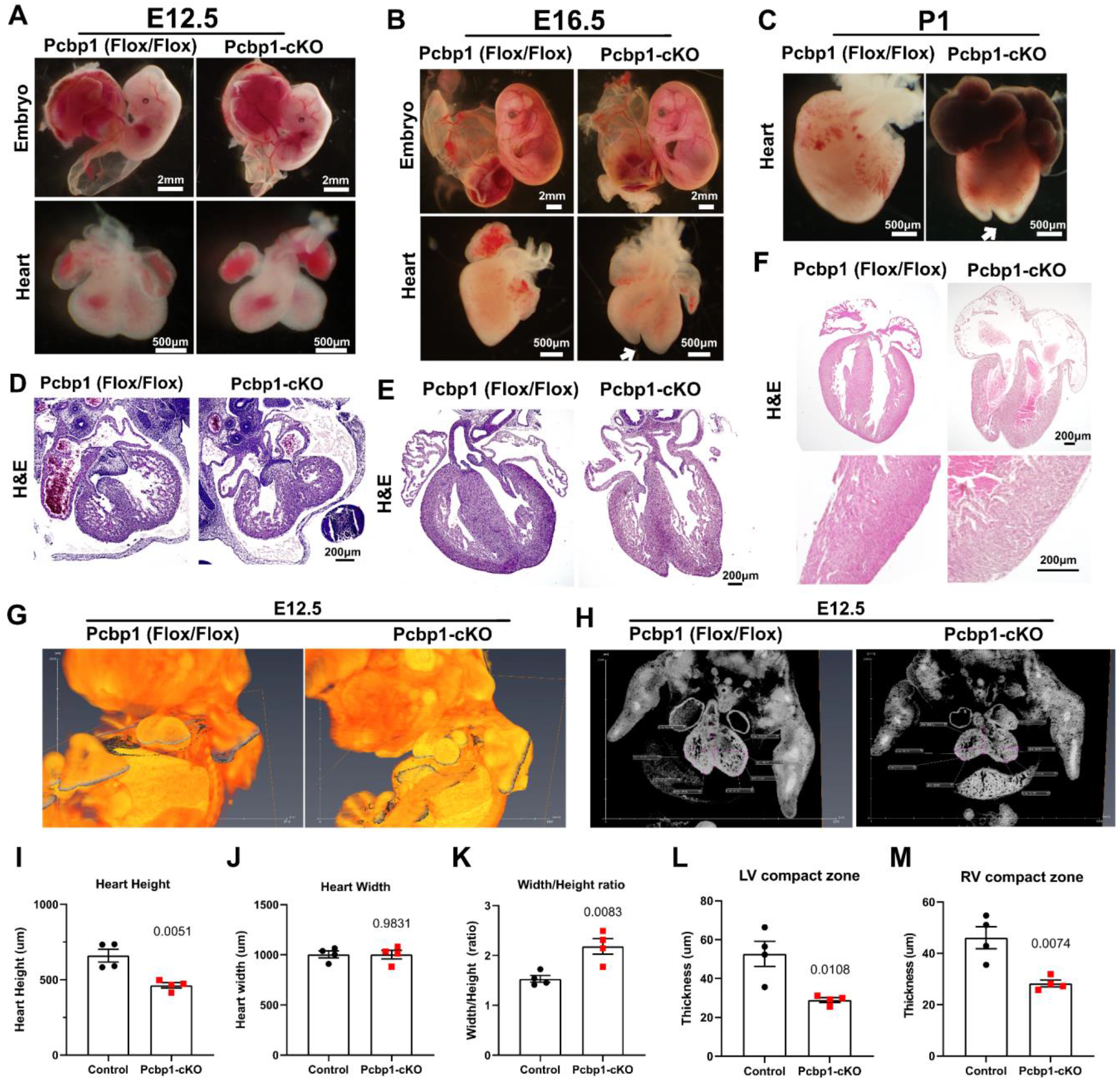
Myocardial deletion of Pcbp1 results in noncompaction cardiomyopathy. Representative gross morphology of the embryo and hearts at E12.5 (A), E16.5 (B) and P1 (C) were shown. White arrow indicates the abnormal apex bifurcation observed at E16.5 and P1. Histological serial sectioning was performed on embryonic hearts of E12.5 (D), E16.5 (E) and P1(F) followed by Hematoxylin and Eosin (H&E) staining (n=7 for each developmental stage). (G) E12.5 embryos with Pcbp1 myocardial knockout and their littermate controls (Pcbp1 Flox/Flox) were processed for high-resolution episcopic microscopy and 3D reconstruction (n=4). 3D rendered volume for representative Pcbp1(Flox/Flox) and Pcbp1-cKOcTNT embryos were displayed. The grey plane indicates the oblique optical sections that coronally slice through the embryonic hearts. (H) The cross-section view from the coronally sliced section were shown. From the coronal plane of each embryonic heart, the height (I), width (J), the width/height ratio (K), as well as the thickness of the compact zone of the left ventricle (L) and right ventricle (M) were measured (n=4, Student’s T-Test p-value shown, data presented as mean ± SEM).

We monitored the Pcbp1-cKO neonatal pups immediately after birth (P0) and found that the majority of pups survived for only 1-2 days postnatally, with 57.2% of the observed Pcbp1-cKO death by P2 (**Supplemental Figure 2F**). To evaluate the neonatal cardiac function, we performed echocardiograms for a cohort of P0, P1, and P2 pups. We found that Pcbp1-cKO pups were born with significantly lower % fractional shortening (%FS), as well as an elevated systolic LVID;s and LV Vol;s, with unchanged LVID;d, LV Vol;d, LVPW;s, and LVPW;d, suggesting a ventricular dilation with systolic dysfunction without ventricular wall thickness change (**Supplemental Table 1**). The cardiac function continued to decrease at P1 and P2, with further reduction in systolic function as indicated by an elevated LVID;s, LVPW;s and LV Vol;s (**Supplemental Tables 2 and 3**). The progressive reduction of neonatal systolic function is consistent with the outcome from most LVNC patients^36^. Together, these data suggest Pcbp1 is essential for ventricular compaction, apex development, and the preservation of neonatal cardiac function.

### Pcbp1 deletion resulted in reduced NOTCH signaling and an elevated unfolded protein response in the heart

To investigate the molecular changes in the absence of Pcbp1, we performed deep RNA-sequencing (RNA-seq) to profile the transcriptome of E16.5 hearts of Pcbp1-cKO animals and Ctnt-Cre negative Pcbp1(Fl/Fl) controls (**Figure 3A**). We identified 154 significantly up-regulated genes and 87 significantly down-regulated genes (|logFC|>0.8, FDR<0.05 cutoff). Hallmark pathway analysis performed using gene set enrichment analysis (GSEA) identified several key gene sets that were dysregulated, such as “Notch signaling” and “Unfolded protein response” (UPR) (**Supplemental Figure 3I**). Ventricular trabeculation and compaction requires proper Notch signaling between the endocardium and myocardium_38,39_, as disruption of the Notch ligand in the endocardium or deletion of Notch effectors in the myocardium leads to hypertrabaculation and non-compaction_39–41_. We found that Notch2, Jag1, and Hey2 were significantly reduced in the Pcbp1 knockout hearts at E12.5 and E16.5 (**Figure 3C-E, Supplemental Figure 3C-D**). Notch activation results in up-regulation of Hey2 in the myocardium during cardiac chamber formation and maturation, and it has been proposed to maintain ventricular identity in the compact myocardium via suppression of atrial genes such as Nppa_42,43_. We found that Nppa expression was highly up-regulated in the Pcbp1-cKO heart in both E12.5 and E16.5 (**Supplementary Figure 3H,** **Figure 3H**). To investigate the spatial distribution of Hey2 and Nppa mRNAs, we performed RNAscope labeling, co-immunostained with sarcomeric myosin (MF20) and Emcn, to identify the compact and trabeculae zones of the myocardium. We found Hey2 mRNA to be largely restricted to the compact myocardium and expressed at a lower level in the thinner compact zone of the Pcbp1-cKO heart (**Figure 3F**). The Nppa mRNA was highly expressed in the Emcn+ trabeculae region, and lower or absent in compacted myocardium in the control E16.5 heart. In contrast, the Nppa mRNA was ectopically expressed at a high level in the compact myocardium in the Pcbp1-cKO heart (**Figure 3G**). These results indicate a critical role for Pcbp1 in mediating Notch signaling and cardiac chamber maturation in the developing heart.

**Figure 3.**
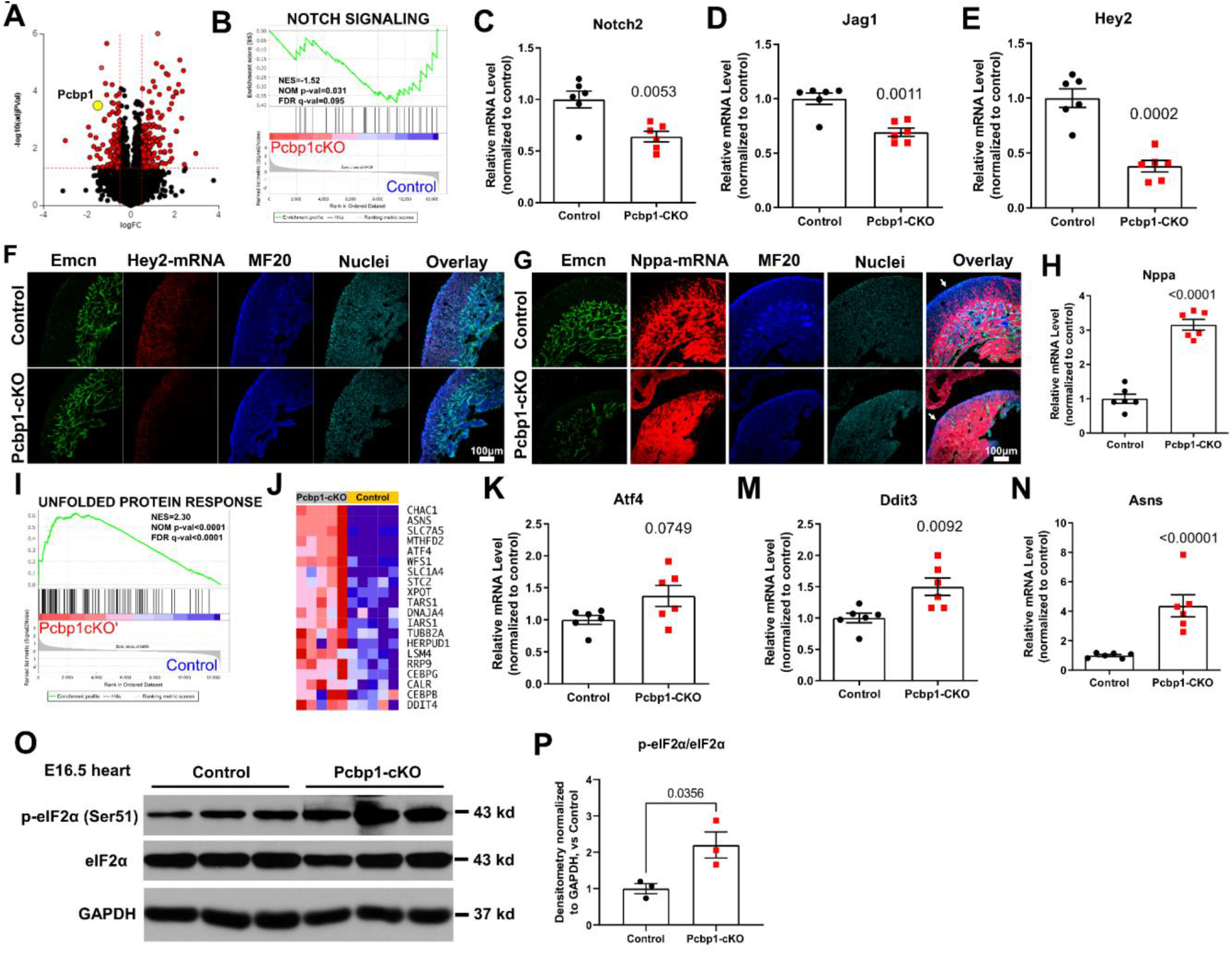
Myocardial deletion of Pcbp1 leads to gene expression dysregulation of NOTCH and UPR pathways. Bulk RNA-seq was performed on E16.5 hearts from Pcbp1-cKOcTNT and littermate controls (n=5). (A) Volcano plot displays the differential gene expression. (B) GSEA analysis was performed with Hallmark gene-set enrichment, and Notch signaling pathway enrichment was shown. Representative myocardial Notch genes were assayed by qRT-PCR on E16.5 hearts (C-E) (n=6, Student’s T-Test p-values shown, data presented as mean ± SEM). (F) RNAscope labeling of Hey2 mRNA and co-immunostaining of Emcn, MF20, and Hoechst 33342 for nuclei (representative images of n=4). (G) RNAscope labeling of Nppa mRNA and co-immunostaining of Emcn, MF20, and Hoechst 33342 for nuclei (representative images of n=4). (H) Nppa transcript was assayed by qRT-PCR on E16.5 hearts (n=6, Student’s T-Test p-values shown, data presented as mean ± SEM). Unfolded Protein Response (UPR) dysregulation with GSEA analysis was shown in (I) and (J). Representative UPR genes, Atf4 (K), Ddit3 (M), and Asns (N) were assayed by qRT-PCR on E16.5 hearts. (n=6, Student’s T-Test p-values, data presented as mean ± SEM). (O) Total and Ser 51 phosphorylation of eIF2α in the E16.5 heart was measured by Western blotting, with GAPDH as a housekeeping control. (P) The p-eIF2α and eIF2α ratio of their respective densitometry measurements normalized to that of GAPDH is shown (n=3, Student’s T-Test p-values, data presented as mean ± SEM).

The unfolded protein response (UPR) is a highly conserved protein quality control system that is activated when cells have impaired folding of nascent proteins in the endoplasmic reticulum_44_, or in the mitochondria during mitochondrial dysfunction_45_. Mitochondrial dysfunction also induces the integrated stress response (ISR), a branch of the UPR mediated by Atf4 and involving the phosphorylation of eukaryotic translation initiation factor 2 alpha (eIF2α) _46–53_. Through GSEA, we found the Pcbp1-cKO hearts had markedly increased UPR-related gene expression (**Figure 3I,J**). UPR/ISR downstream genes such as Ddit3, amino acid nutrient stress gene Asns_54_, and Atf4 were elevated in E12.5 and E16.5 hearts (**Supplemental Figure 3E-G and** **Figure 3K-N**), and levels of the Serine 51 phosphorylation of eIF2α (p-eIF2α) were more than two-fold higher in the E16.5 heart (**Figure 3O, P**). These data suggest that UPR/ISR-induced stress is prominent in the Pcbp1-cKO heart.

### Disrupted cardiomyocyte maturation trajectory in Pcbp1 mutant hearts

Since the reduction of Hey2 mRNA and elevated Nppa mRNA in the compact myocardium indicated a shift of signaling events related to a delay in maturation, we sought to assess the maturation state of the Pcbp1-cKO heart using an unbiased approach. We took existing bulk RNA-seq datasets of E10.5, E11.5 (early) and E15.5, E16.5 (late) heart ventricles (ENCSR049UJU, ENCSR691OPQ, ENCSR020DGG, ENCSR597UZW)_55_, and used the differentially expressed leading edge genes (**Figure 4C**) from comparisons of the early and late heart development stages to construct gene sets that correspond to up-regulated and down-regulated genes in E16.5 hearts. We then ran our Pcbp1-cKO heart transcriptome against these gene sets using GSEA and found that Pcbp1 deficiency in the myocardium caused a positive enrichment of genes that are normally down-regulated in E16.5 hearts (**Figure 4D**), and a negative enrichment of genes that are normally up-regulated in E16.5 hearts (**Figure 4E**). These data demonstrated a clear transcriptomic shift and dysregulation in the developmental trajectory of ventricular maturation associated with Pcbp1 deficiency.

We then validated these findings using RNAscope *in situ* hybridization to label chamber maturation-related expression in E16.5 hearts. Myl4, Myl7, and Bmp10 has been shown by *in situ* hybridization to express ubiquitously in the atrium and ventricles during early cardiogenesis, and their expressions become atrial-restricted in late cardiac development_42,43_. We found that in the control E16.5 heart, Myl4, Myl7 (**Figure 4F**), and Bmp10 mRNAs (**Figure 4G**) were restricted to the atrium in the wild-type control E16.5 heart, consistent with previous studies_42,43_. In the absence of Pcbp1, we detected ectopic expression of Myl4, Myl7, and Bmp10 mRNAs in the myocardium and trabeculae zones (**Figure 4F and G**), suggesting that Pcbp1 deficiency results in ventricular developmental delay.

**Figure 4.**
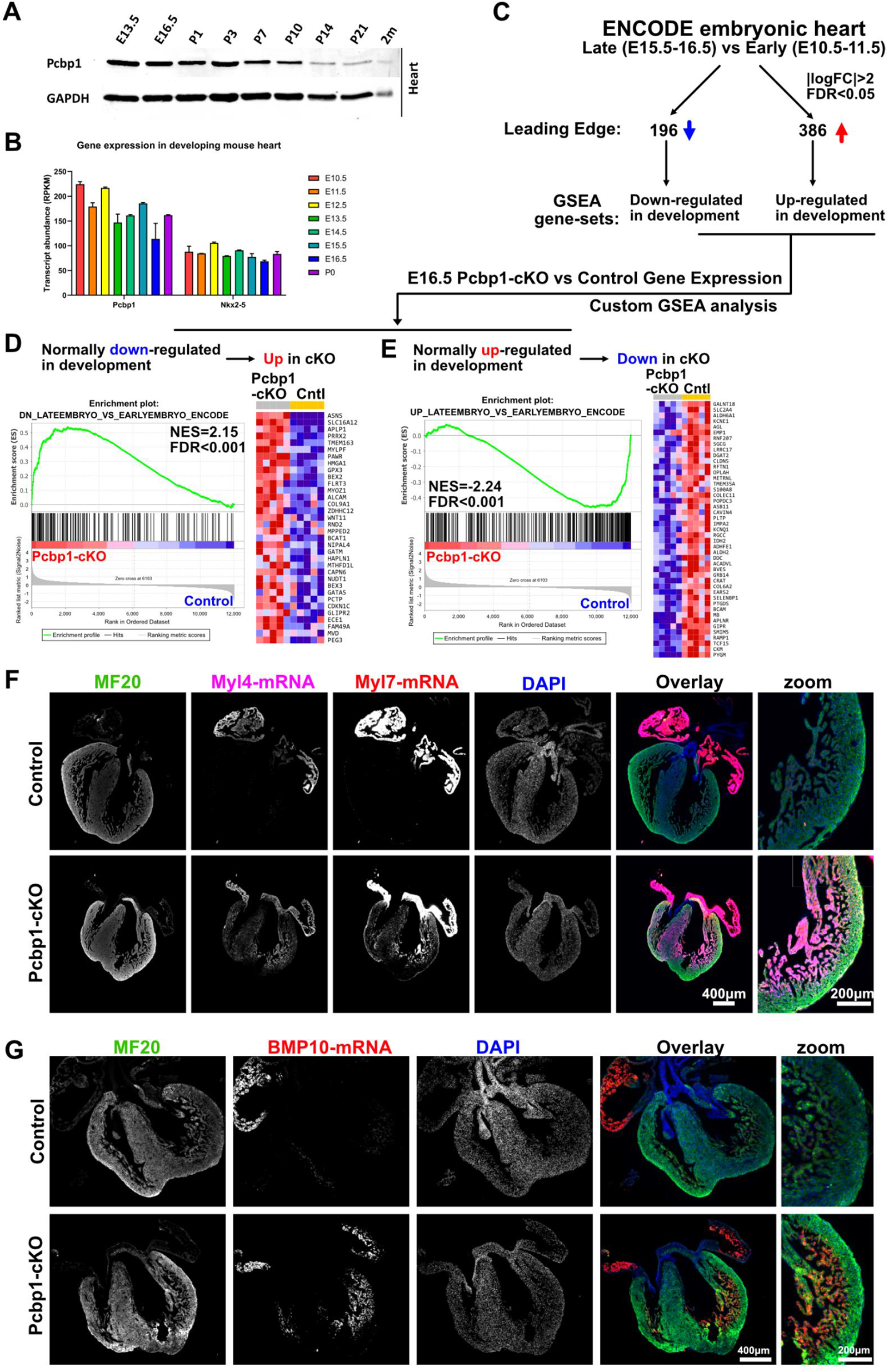
Myocardial Pcbp1 deletion caused a delay in ventricular maturation during development. (A) The heart ventricles during embryonic development and postnatal time points were harvested and Western blotting was performed to measure Pcbp1 protein expression, with GAPDH as a housekeeping control (n=3). (B) Transcript level of Pcbp1 was assessed from developmental heart tissues from ENCODE, the levels of Pcbp1 and Nkx2.5 were shown (n=2 per timepoint). (C) Using the ENCODE embryonic heart RNA-seq dataset, we assigned E15.5 and E16.5 RNA-seq data as “Late” timepoint group, the E10.5 and E11.5 RNA-seq data as “Early” timepoint group (n=4 per group), and we compared the Early and Late group and extracted the leaging edge of highly significant differentially expressed genes, and constructed two custom GSEA gene-sets for up-regulation (“UP_LATEEMBRYO_VS_EARLYEMBRYO_ENCODE”) and down-regulation (“DN_LATEEMBRYO_VS_EARLYEMBRYO_ENCODE”) that summarize the developmentally differentially regulated genes in the heart. (D-E) These developmentally differentially regulated gene-sets were subsequently applied to the RNA-seq data from Pcbp1-cKO compared with control. The Normalized Enrichment Score (NES) and FDR values were reported, and the heatmaps of the leading edge of the differentially regulated genes within each GSEA analysis were shown. (F) RNAscope labeling of Myl4 and Myl7 mRNAs with co-immunofluorescence staining of MF20 and DAPI were shown (n=3). (G) BMP10 mRNA was also examined by RNAscope with co-immunoflorecence staining of MF20 and DAPI (n=3).

### PCBP1 regulates alternative splicing of AARS2 in the heart

Since PCBP1 has been shown to be capable of regulating alternative splicing (AS) of targets such as CD44 in HepG2 cells and Runx1, EpoR, Epb41, and Epb49 in primary erythrocytes _12,22,56,57_, we sought to define the alternative splicing landscape regulated by PCBP1 in the heart. Using rMATs (v4.0.2) _58,59_, we assessed the levels of alternative exon inclusion, expressed as “percent spliced in” (Psi) and compared the Pcbp1-cKO hearts with control hearts. Of both annotated and novel alternative splicing events detected, we found 542 alternatively spliced genes that can be classified into one of the five previously defined major classes of splicing events: A3SS (alternative 3’ splice site), A5SS (alternative 5’splice site), SE (skipped exon), RI (retained intron), and MXE (mutually exclusive exons) (**Supplemental Figure 4D)**. We found 341 SE events in hearts with Pcbp1 deficiency, constituting 62.92% of the total detected AS events (**Figure 5A, Supplemental Figure 4E and F**). We found 84 RI events which comprise 15.5% of the AS events, and the rest of the categories account for less than 8% each (**Figure 5A, Supplemental Figure 4G-J**). It is noteworthy that the AS events we found in the heart caused by Pcbp1 deficiency have minimal overlap with the previous described AS events in HepG2 cells or erythrocytes, indicating a cell-type specificity of AS regulation. Together, this suggests that the dysregulation of AS caused by Pcbp1 deficiency in the heart is comprised predominantly of SE events.

**Figure 5.**
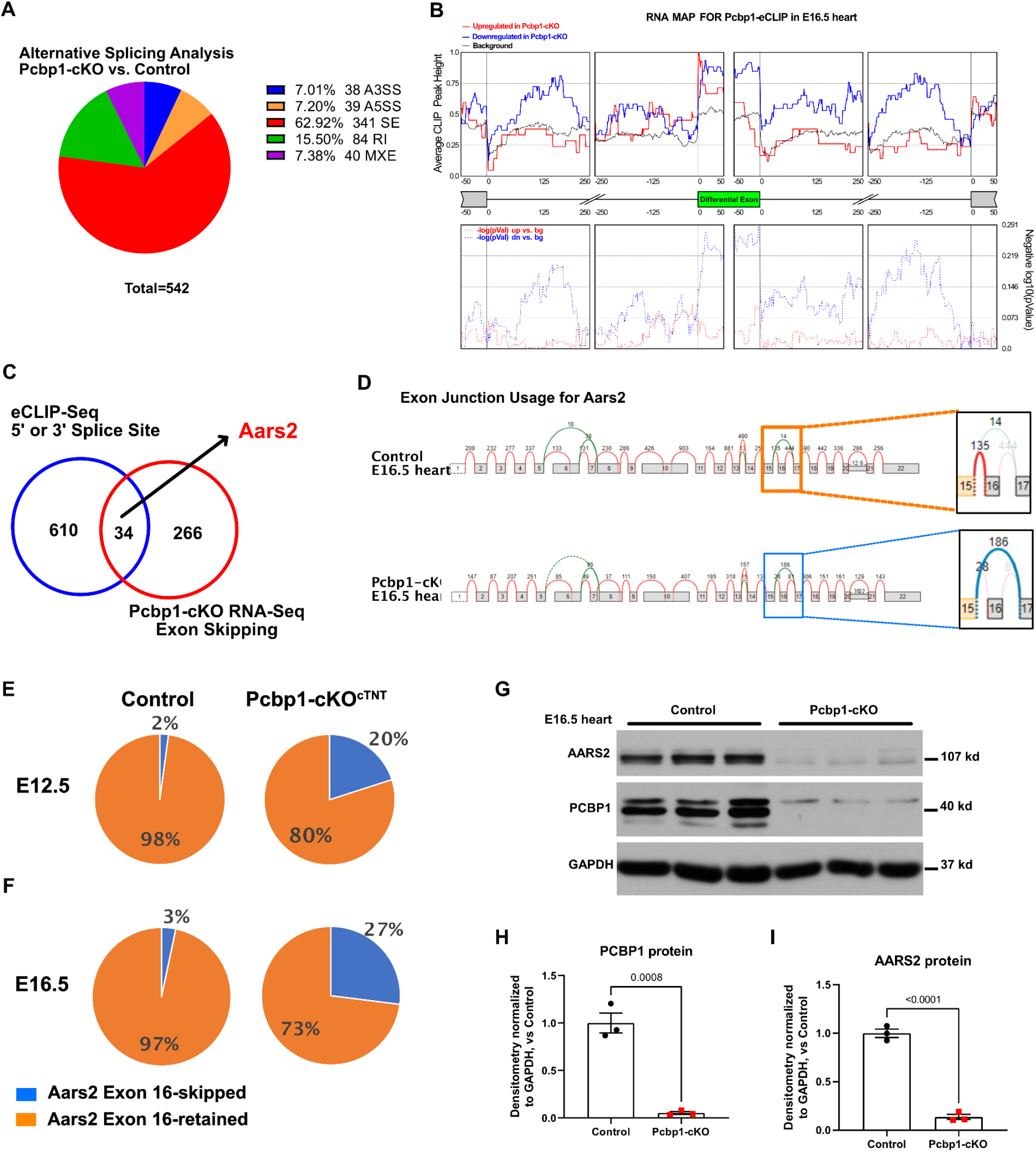
Deletion of Pcbp1 caused aberrant alternative splicing of Aars2 through interaction with intronic splice site. (A) Alternative splicing analysis performed on the E16.5 Pcbp1-cKO transcriptome identified 542 alternative splicing events. Among those, the skipped exon (SE) events were over-represented and included 62.97% of total events. (B) By integrating of the PCBP1 eCLIP-seq peaks and alternative spliced transcripts from the transcriptome of Pcbp1-cKO RNA-seq, we identified a significant enrichment of PCBP1 eCLIP-seq signal at the 5’ and 3’ flanking regions of the skipped exon from the Pcbp1-cKO transcriptome. (C) We focused on gene loci with PCBP1 eCLIP peaks at the 5’ or 3’ splice sites, and genes with exon-skipping induced by Pcbp1-cKO, and found that Aars2, a CHD casual gene, is among the exon-skipped genes by Pcbp1-cKO with eCLIP binding. (D) The alternative splicing of exon 16 of Aars2 is shown, the zoom-in view highlights the junctional reads from exon 15 were predominantly skipped to exon 17 in the Pcbp1-cKO heart. (E-F) The expression of the exon-16 skipped and exon-16 retained transcripts were quantified using absolute qRT-PCR with respective amplicons and standard curves. We found that at both E12.5 (F) and E16.5 (G), Pcbp1-cKOcTNT has about 10-fold more exon-16 skipped transcripts (n=6). (G) The protein levels of AARS2 and PCBP1 in the Pcbp1-cKO and control E16.5 hearts were measured by Western blotting, with GAPDH as a housekeeping control. (P) The densitometry measurements normalized to that of GAPDH for AARS2 (H) and PCBP1 levels (I) were shown. (n=3, Student’s T-Test p-values are shown, data presented as mean ± SEM).

We sought to test whether the SE regulation by Pcbp1 can be generalized to the transcriptome level. Leveraging the Pcbp1 eCLIP binding and the SE regulated loci identified from the Pcbp1-cKO transcriptome in the same developmental stage of E16.5, we generated an RNA map of Pcbp1-interacting transcripts with AS data using rMAPs (v2.2.0) _60,61_ (**Figure 5B**). Through analysis of Pcbp1’s binding location, exon loci that are down-regulated (exon-skipped) in the Pcbp1-cKO had enrichment of Pcbp1 eCLIP peaks at the 5’ and 3’ flanking intronic regions, and no significant peak enrichment in loci with up-regulated exon (exon-inclusion). Importantly, no significant enrichment of Pcbp1 eCLIP signal can be identified from other alternative splicing categories such as A3SS, A5SS, RI, or MXE (**Supplemental Figure 5**), which further suggests a direct regulatory function for Pcbp1 in stabilizing skipped exons in the developing heart.

Since most of the alternative splicing events identified in the Pcbp1-deficient heart are skipped exons, we focused on genes that exhibit exon-skipping in the Pcbp1-cKO with Pcbp1 mRNA binding at the 5’ or 3’ splice site. By comparing the genes identified with exon-skipping events in the Pcbp1-cKO, with the genes harboring Pcbp1 eCLIP peaks located near the splice sites, we narrowed the list down to 34 genes that have both features (**Figure 5C**). We found that AARS2, a gene that we identified to interact with PCBP1 near exon 16 (**Figure 1**), displayed exon-16 skipping. Strikingly, this exon-skipping has not been previously identified or annotated in the transcriptome, and its exon junction usage is being robustly altered in the absence of Pcbp1 (**Figure 5D**). To quantitatively evaluate the splicing change, we designed a quantitative qPCR approach to detect and quantify the exon-16 skipped or exon-16 retained transcript (**Supplemental Figure 3M-P**), similar to previously described quantitative methods for exon-skipping and inclusion _62–64_. We found that the Aars2 exon-16 skipping can be detected as early as E12.5 in Pcbp1-cKO to comprise 20% of the total Aars2 transcript (**Figure 5E**), and the proportion of the exon-skipping transcript increased further at E16.5 to 27% (**Figure 5F**). Notably, the Aars2 exon-16 skipped transcripts were almost absent in the E12.5 and E16.5 control hearts, where only 2-3% of the Aars2 transcripts exhibited exon-16 skipping. The exon-16 skipping of Aars2 causes a downstream frameshift that produces a premature termination signal (**Supplemental Table 4**), and it will likely affect its transcript stability and protein truncation. The mRNA level for Aars2 as assessed by the non-spliced exon-14 to exon-15 region was reduced by 59% (**Supplementary Figure 3L**), indicating the overall transcript level for Aars2 is also decreased, potentially through nonsense-mediated decay mechanism elicited to eliminate the mutant transcripts. To evaluate the protein level of AARS2 in Pcbp1-cKO, we isolated protein from E16.5 ventricles from control and Pcbp1-cKO embryos and performed Western blotting (**Figure 5G**). As expected, Pcbp1 protein levels were significantly diminished in the Pcbp1-cKO heart and we detected a striking decrease of AARS2 protein level as well, with only 13.7% of AARS2 remaining as assessed by densitometry (**Figure 5H,I**). Overall, this evidence suggests a robust regulatory role for PCBP1 in regulating AARS2 exon-16 skipping and its protein level in the developing heart.

### Aars2 exon-16 deletion causes ventricular non-compaction and apex malformation

To directly test the consequence and effects of Aars2 exon-16 skipping, we used the homology recombination-directed CRISPR/Cas9 strategy to insert loxP sites flanking exon-16 of Aars2 using a single donor template (**Figure 6A, and Supplemental Figure 6A,B**) together with two different combinations of gRNA pairs with PAM sites located in the flanking introns of exon-16. We screened the founder progenies by genomic PCR followed by Sanger Sequencing to verify the location of the insertion and deletion and successfully recovered an allele with the correct LoxP-sites inserted, as well as mutants that harbored deletions of various sizes and at different locations (**Supplemental Figure 6C-E**). After back-crossing three generations to C57BL/6 to minimize effects from potential off-target editing, we chose three deletion alleles on or near the exon-16 of Aars2. Aars2-DEL248 harbors a 251 bp deletion of exon-16, with most of the intronic element preserved, and is predicted to frame-shift the rest of the transcript (**Supplemental Figure 7A**). Aars2-DEL-IF harbors a 320 bp deletion that is flush to the 3’ end of exon-15 and removes one extra nucleotide on the 5’ end of the exon-17; as a consequence, this Aars2-DEL-IF is predicted to remove exon-16 while keeping exon-17 onwards in-frame (**Supplemental Figure 7B**). Aars2-In70 harbors a 43 bp deletion at the intron between exon-15 and exon-16; interestingly, the deleted region contains a consensus Pcbp1 motif (**Supplemental Figure 7C**). All three heterozygous mutants are viable and fertile. We then crossed these germline heterozygous mutants independently to homozygosity; however, we could not obtain viable progenies at and after E8.75, suggesting that all three homozygous mutants resulted in early embryonic lethality (**Supplemental Figure 7**). The fact that both in-frame Aars2-DEL-IF and frame-shift Aars2-DEL248 deletion mutants resulted in the similar early embryonic lethality suggests that Aars2 exon-16, regardless of downstream frameshift or its intronic genomic sequence, is essential for early embryonic survival.

**Figure 6.**
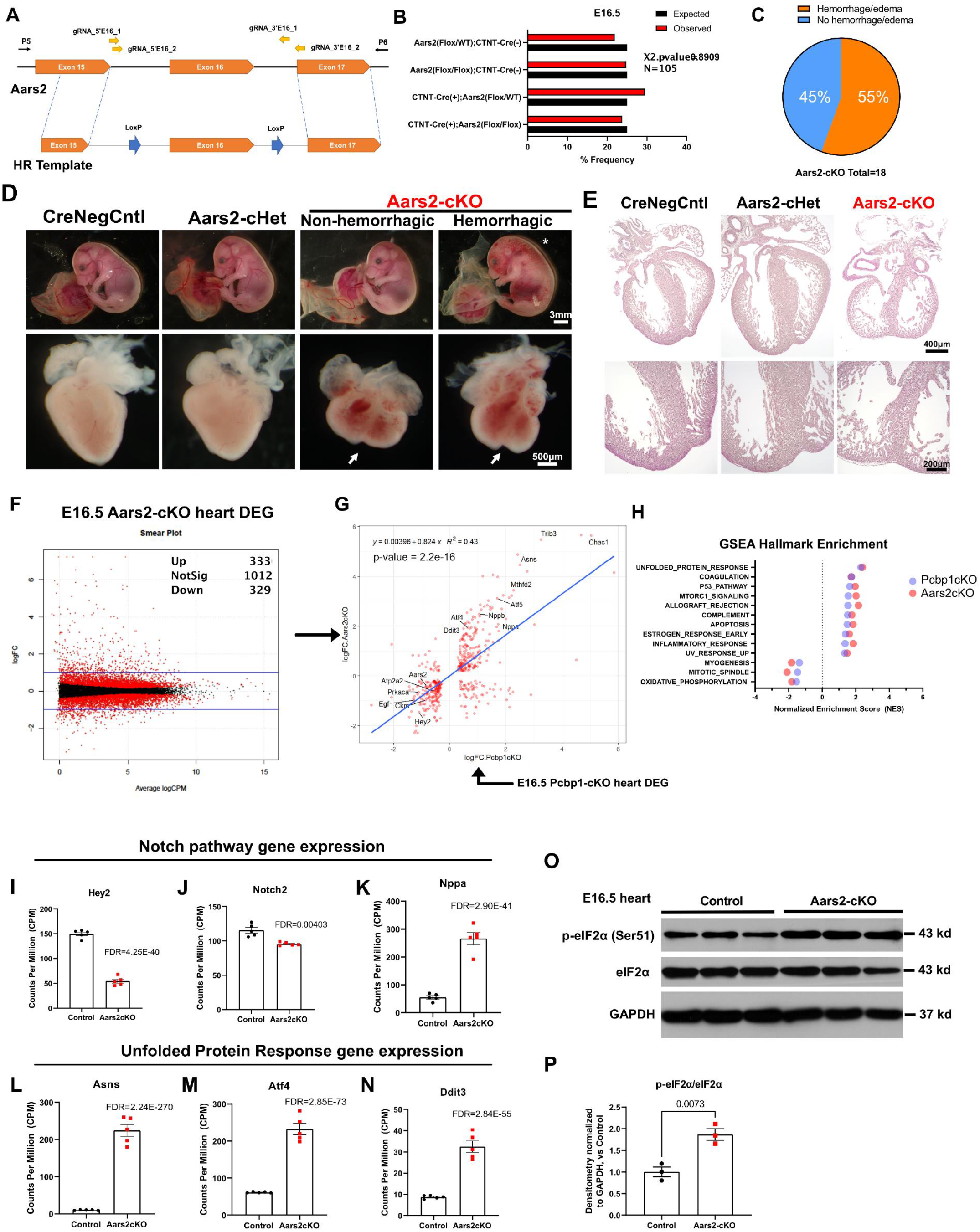
Myocardial deletion of Aars2 exon-16 causes similar myocardial non-compaction and disruption of gene expressions, phenocopying the Pcbp1-cKO heart. (A) A new mouse line with loxP sites flanking exon-16 of Aars2 was generated with CRISPR/Cas9 mediated knock-in strategy with the indicated DNA template, and subsequently we bred this line with cTNT-Cre mice to generate myocardial deletion of Aars2 exon-16. (B) Expected and observed frequency at E16.5 for the cross was shown (C) At E16.5, Aars2-cKO embryo has 55% penetrance of displaying hemorrhage and edema. (D) Gross embryonic and cardiac morphology of Aars2-cKO (D) and H&E staining of heart sections (E) indicated a severe ventricular non-compaction and apex malformation (white arrow). (F) We collected E16.5 hearts from Aars2-cKO and controls (n=5) and performed bulk RNA-seq, with differentially expressed genes (DEGs) shown in a smear plot (red dots = DEGs with FDR < 0.05). (G) Linear regression analysis was performed on the DEGs of Pcbp1-cKO (x-axis) and Aars2-cKO (y-axis) (FDR = 0.05, log2FC = 0.5), with linear relationship between these two variables shown with high significance. (H) GSEA analysis with the Hallmark gene-set showed a high concordance in dysregulated pathways between Pcbp1-cKO and Aars2-cKO. Lastly, we assayed myocardial Notch pathway genes (I, J, K), as well as UPR genes (L, M, N) of E16.5 hearts from Aars2-cKO and littermate controls (n=5, Student’s T-Test p-values, data presented as mean ± SEM). (O) Total and Ser 51 phosphorylation of eIF2α in the E16.5 heart was measured by Western blotting, with GAPDH served as housekeeping control. (P) The p-eIF2α and eIF2α ratio of their respective densitometry measurements normalized to that of GAPDH was shown (n=3, Student’s T-Test p-values shown, data presented as mean ± SEM).

The Aars2 E16 loxP (Aars2-Flox) mice (**Figure 6A, Supplemental Figure 6F-H**) were viable and fertile when bred to homozygosity and were used to generate a cardiomyocyte-specific deletion of Aars2 exon-16 by crossing with Ctnt-Cre mice (Aars2-cKO). We could not recover any viable Aars2-cKO at P1, although we observed the expected Mendelian ratio at E16.5 (**Figure 6B**). Of all the Aars2-cKO embryos recovered at E16.5, 55% of them displayed gross hemorrhage/edema throughout the embryo, while 45% of them were normal (**Figure 6C**). The incomplete penetrance of the hemorrhage and edema phenotype could be a secondary consequence of deteriorating cardiac function caused by the Aars2 exon-16 deletion. The Aars2-cKO E16.5 heart had a shorter statue and apex malformation, similar to that of Pcbp1-cKO (**Figure 6D**). Histological sectioning of the heart followed by H&E staining revealed hyper-trabeculation and severe thinning of the myocardium (**Figure 6E**), indicating a similar ventricular non-compaction phenotype observed in the Pcbp1-cKO.

### Both Pcbp1 and Aars2 participate in the regulation of oxidative phosphorylation in cardiomyopathy

Since the phenotype for Aars2-cKO showed many attributes similar to those of the Pcbp1-cKO, we hypothesized that the gene expression alterations from these two models would share similar dysregulation. We performed bulk RNA-seq from Aars2-cKO hearts and their controls at the same embryonic stage (E16.5) and found significantly more genes dysregulated in Aars2-cKO hearts compared with Pcbp1-cKO hearts (**Figure 6F**). This observation is consistent with a more severe phenotype observed in the Aars2-cKO embryo. We performed linear regression with differentially expressed genes of Aars2-cKO and Pcbp1-cKO, and found that the dysregulated genes from Aars2-cKO positively correlated with those from Pcbp1-cKO, with an R_2_ value of 0.43 (**Figure 6G**). Most of the dysregulated genes located on the top-right or bottom left quadrant, suggesting that these sets of genes are concordantly dysregulated. Indeed, we found 42.4% and 43.8% of the genes either up-or down-regulated in Pcbp1-cKO were also dysregulated in the Aars2-cKO (**Figure 5C and D**). We performed GSEA with the Hallmark gene sets and found remarkable similarity in the gene sets that were enriched, as well as the normalized enrichment score that aligned in the same direction and in similar magnitude. Together, these data suggest remarkably similar gene expression and pathway alterations in the Pcbp1-cKO and Aars2-cKO models, suggesting that Pcbp1 and Aars2 may be crucial for regulating the same pathways during heart development.

To assess the gene expression changes for Notch and the UPR/ISR, we performed qRT-PCR on RNA isolated from E16.5 Aars2-cKO and littermate control hearts. We found that compact myocardium enriched genes Hey2 and Notch2 were significantly decreased, and the atrial enriched gene Nppa was elevated (**Figure 6I-K**). The UPR/ISR genes Atf4, Asns, and Ddit3 were all significantly up-regulated in the Aars2-cKO heart (**Figure 6L-M**). Importantly, we also detected a significant increase in p-eIF2α in the Aars2-cKO (**Figure 6O, P**), indicating that the UPR/ISR-induced stress is also prominent in the Aars2-cKO hearts. We observed an overall up-regulation of aminoacyl-tRNA transferase (ARS) in both Pcbp1-cKO and Aars2-cKO (**Supplemental Figure 8A,B**): specifically, 11 of the cytosolic ARS transcripts were concordantly up-regulated (**Supplemental Figure 8C**). The activation of cytosolic ARS was previously attributed to transcriptional activation through Atf4_65_. Together, this suggests a unified shift in UPR/ISR stress response and ventricular maturation in both the Pcbp1-cKO and Aars2-cKO heart.

One of the hallmarks for the AARS2 mutation in congenital cardiomyopathy is the deficiency of the oxidative phosphorylation complexes (OXPHOS)_6,25_, and patient samples have overall reduction of complex I (C-I) protein and variable reduction in other OXPHOS complexes. Therefore, we sought to examine the OXPHOS complexes in the Pcbp1-cKO and Aars2-cKO embryonic hearts. We found the transcriptomes of both Pcbp1-cKO and Aars2-cKO hearts had a significant down-regulation of OXPHOS-related gene sets (**Figure 7A,B**). Through examining the protein lysates from E16.5 hearts, we found protein levels of NDUFB8 (C-I) and SDHB (C-II) were consistently abolished in both the Pcbp1-cKO and Aars2-cKO heart. Additionally, the UQCRC2 (C-III) and vATP5A (C-V) levels were significantly down-regulated in Aars2-cKO but not in the Pcbp1-cKO heart. Overall, we found that OXPHOS deficiency is present in both the Aars2-cKO and Pcbp1-cKO, consistent with observations in patients.

**Figure 7.**
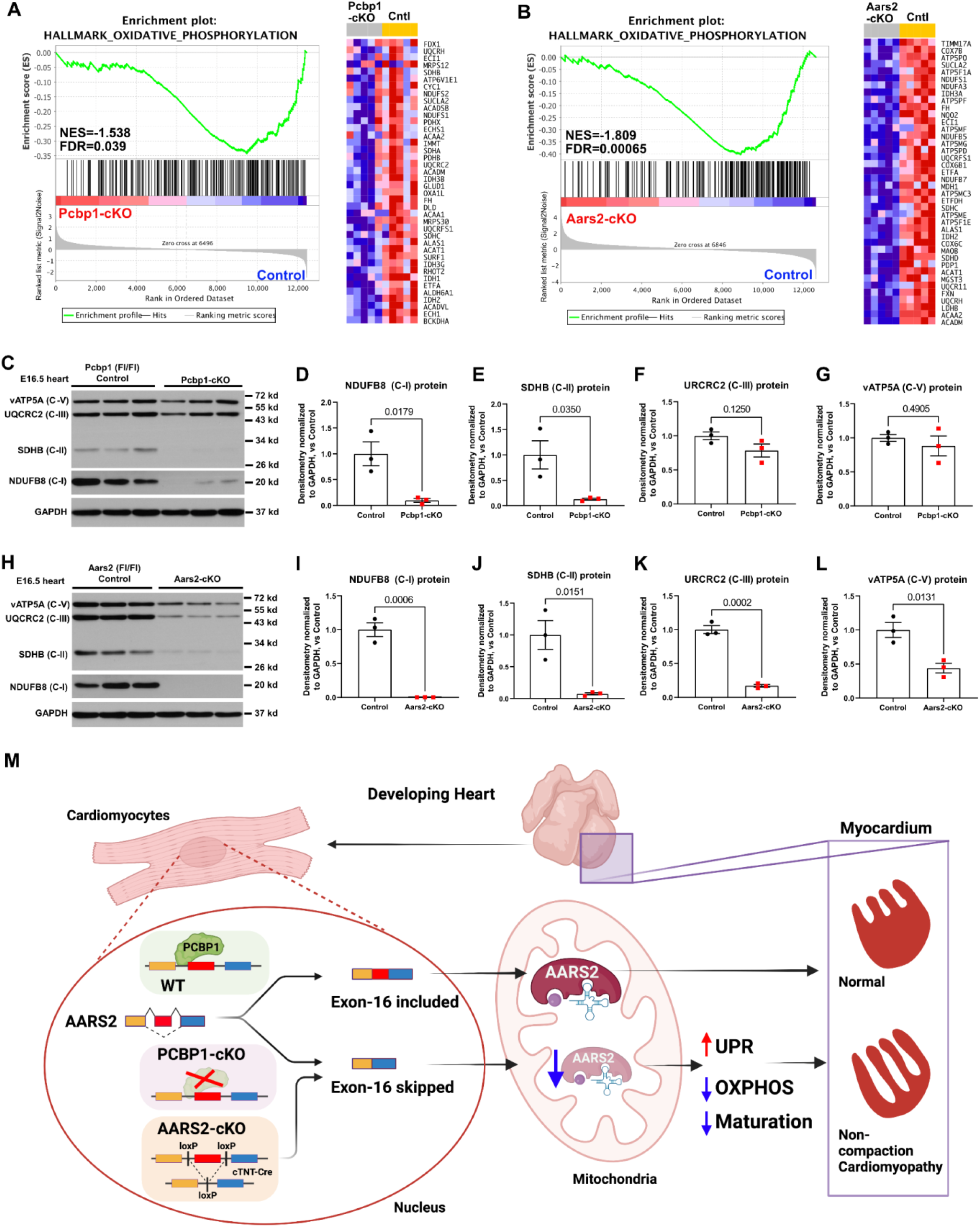
Myocardial deletion of Pcbp1 or Aars2 exon-16 both lead to decreased protein levels of oxidative phosphorylation complexes. GSEA enrichment analysis for the Hallmark oxidative phosphorylation gene-set enrichment for Pcbp1-cKO (A) and Aars2-cKO (B) are shown accordingly. The protein level of key components from the oxidative phosphorylation complexes, such as vATP5A, UQCRC2, SDHB, and NDUFB8 were measured by Western blotting, with GAPDH as housekeeping control, for Pcbp1-cKO and control (C), as well as Aars2-cKO and control (H) (n=3). The densitometry for the protein levels were normalized to GAPDH and shown as indicated for Pcbp1-cKO (D-G) and Aars2-cKO (I-L) (n=3, Student’s T-Test p-values shown, data presented as mean ± SEM). (M) Summary schematic showing that we identified PCBP1 as a novel regulator for AARS2 alternative splicing, which is essential for cardiac ventricular morphogenesis. Myocardial deletion of Pcbp1 causes exon-16 skipping of Aars2 through a direct interaction. Myocardial deletion of Aars2 exon-16 phenocopies Pcbp1-cKO with oxidative phosphorylation deficiency, contributing to the non-compaction cardiomyopathy phenotype.

Previous studies had shown that direct perturbation of OXPHOS in the embryonic heart can lead to severe heart defects and lethality_66–71_. Our genetic approach following a congenital mitochondrial cardiomyopathy causal gene, AARS2, has uncovered a novel mechanism that involves the regulation of PCBP1 on the alternative splicing and functions of AARS2 during cardiac development.

## DISCUSSION

In this study, we report that PCBP1 plays a pivotal role in regulating gene expression and alternative splicing of AARS2 in cardiomyocytes during heart development. We show that cardiac-specific loss of Pcbp1 during development caused myocardial non-compaction, apex malformation, and perinatal lethality, with reduced Notch signaling and activated UPR/ISR. Through a multimodal approach with the Pcbp1-cKO transcriptome and Pcbp1-RNA interactome in the embryonic heart, we identified Aars2, a congenital mitochondrial cardiomyopathy causal gene_6,72_, as a novel alternative splicing target of Pcbp1. Specifically, Pcbp1 interacts with Aars2 mRNA regions flanking exon-16 and the absence of Pcbp1 in cardiomyocytes causes an accumulation of an aberrant exon-16 skipping isoform of Aars2. Through generation of a novel transgenic mouse harboring loxP flanking Aars2 exon-16, we conditionally removed this exon in cardiomyocytes during heart development and uncovered a remarkably similar phenotype in Aars2-deficient hearts that displayed myocardial non-compaction and apex malformation. Further transcriptome analysis uncovered a strikingly concordant dysregulation of genes shared between the Pcbp1-deficient heart and the Aars2-deficient heart at E16.5. Our study revealed a novel function for Pcbp1 in regulating cardiac development, in part by maintaining the exon-16 inclusion of Aars2. The identification of Pcbp1 as a critical player in non-compaction cardiomyopathy and ventricular morphogenesis provides important insights into the requirement of RBP function for proper formation of the heart through the regulation of Aars2 alternative splicing.

### Roles of Pcbp1 in heart development

The increased complexity of vertebrates compared to invertebrate organisms is believed to be due in large part to the expansion of the transcriptome, without a proportional expansion of the genome_73_; this expansion is due to an increase in pre-mRNA processing and alternative splicing_74_. Consistent with this notion, several RBP families that are critical for the regulation of alternative splicing are expanded in vertebrates relative to invertebrates_75_. RBPs have been shown to regulate cardiac development in multiple stages. A few of the RBPs, such as Rbm20 and Rbfox2, have been linked to human cardiomyopathy and heart development_76–80_. Developmental studies of many of the RBP families, such as the serine/arginine-rich (SR) protein family proteins, had heavily relied on mouse genetic studies due to their requirement for early development prior to heart formation, and many of these RBPs are highly expressed during cardiac development but also ubiquitously expressed in multiple cell types_2_. Pcbp1 is robustly expressed in the heart throughout cardiac development, at developmental stages that involve orchestrated changes in gene expression that transition cardiomyocytes from an immature to matured state. We and others previously found that germline knockout of Pcbp1 results in preimplantation lethality that occurs well before heart formation, suggesting its requirement in the gastrulation stage of early embryonic development_20,21_. A nonsense single nucleotide variant of human PCBP1, c.550C>T, was reported in the ClinVar Database of National Center for Biotechnology Information (NCBI), which changed glutamate in position 184 into a premature termination codon (Gln184Ter), VCV000208707.2 _81_. This variant is predicted to be damaging and likely pathogenic, and the clinical features observed in the individuals with this variant include global developmental delay, proportional short statue, Tetralogy of Fallot, decreased muscle mass, etc., suggesting that the function of PCBP1 is linked to human congenital disease conditions.

In this study, we found that cardiomyocyte-specific knockout of Pcbp1 caused perinatal lethality, non-compaction cardiomyopathy, and defects in ventricular apex formation, highlighting the essential role for Pcbp1 in cardiac development. Though the casual relationship between these genes and LVNC has been debated_82_, Notch pathway dysregulation has been implicated in LVNC in human and mouse models _38–40,42,43,83_. Interestingly, in the context of LVNC, patients with LVNC or mouse models with cardiac-specific deletion of Hey2 – which results in LVNC – often can survive after birth and do not typically display juvenile lethality_42,84_. Therefore, the neonatal lethality of Pcbp1-cKO may be caused by a combined effect of other pathways, such as UPR/ISR or mis-splicing of key regulators. We found that in hearts with Pcbp1 deficiency, UPR/ISR was highly up-regulated at E16.5, with key UPR/ISR genes up-regulated prior to the onset of ventricular compactions at E12.5. Intriguingly, a recent report identified PRDM16 in the myocardium as a critical factor for LVNC in mice_85_, where they generated a mouse model with myocardial deletion of Prdm16 using Ctnt-Cre (Prdm16-cKO) with a robust LVNC phenotype. They observed a consistent down-regulation of Hey2 in the Prdm16-cKO, but no UPR/ISR activation was reported in their bulk RNA-seq data. Together, this suggests that the UPR/ISR activation observed in Pcbp1-cKO may be independent of the dysregulation of the myocardial Notch pathway.

### Alternative splicing of Aars2 causes ventricular non-compaction and perinatal lethality

Currently, AARS2-linked infantile congenital cardiomyopathy has been associated with a recurrent pathogenic p.Arg592Trp (c1774C>T) mutation located in the editing domain of AARS2 that is either homozygous or compound heterozygous in patients_25_. The AARS2 editing domain is required for the deacylation of mischarged tRNAs, due to the inability of the aminoacylation domain to discriminate alanine from serine and glycine_86,87_. This editing mechanism could avoid misincorporation of serine or glycine at the alanine codon during protein synthesis, therefore preventing protein mistranslation. It has been previously proposed that the AARS2-linked cardiomyopathy mutations, which frequently occur within the editing domain, severely compromise the AARS2 aminoacylation activity, whereas partial activity is retained by the mutations found in leukodystrophy patients.

Intriguingly, we identified AARS2 aberrant alternative splicing in the Pcbp1-cKO heart, where the exon-16 encoded part of the AARS2 editing domain is skipped. The direct regulatory role of Pcbp1 in stabilizing this alternative splicing is further supported by our eCLIP-seq and RIP experiments. The loss of Pcbp1 in the heart induces predominantly exon-skipping events, suggesting that the Pcbp1-mRNA interaction may stabilize splicing by suppressing exon-skipping. It is likely that Pcbp1 functions as part of the spliceosome or ribonuclear complexes_88_, and future studies will be directed at identifying other ribonuclear complexes that are involved within this regulatory axis.

Our novel conditional allele of Aars2, in combination with cTNT-Cre, enabled us to remove the Aars2 exon-16 specifically in cardiomyocytes during development. Aars2-cKO mice are embryonically lethal and display LVNC, similar to the Pcbp1-cKO hearts. We found remarkable similarity of the transcriptome of Aars2-cKO and Pcbp1-cKO hearts and that both display OXPHOS deficiency, suggesting they are located within the same regulatory axis. Aars2-cKO was engineered to delete the majority of the exon-16; in contrast, in the Pcbp1-cKO, the exon-16 skipping on Aars2 transcript constitutes only 20-30% of the total transcript, although the majority of the Aars2 protein is depleted. One potential explanation is non-sense mediated decay that has been predicted to deplete the targeted transcript, thereby having a dominant negative effect on its expression_89_. It is conceivable that in addition to the regulation of Aars2 alternative splicing, which explains the majority of the transcriptome changes in Pcbp1-cKO, Pcbp1 might also regulate Aars2-independent gene expression and thereby contribute to the phenotype. The fact that Aars2-cKO has a more severe phenotype in comparison to Pcbp1-cKO, would suggest the Aars2-independent programs regulated by Pcbp1 might oppose the damaging effects elicited by Aars2 depletion alone. It would be of interest to further delineate the additional regulatory pathways that branch from the PCBP1-AARS2 axis.

Since the Tfam transcript, as well as mitochondrial encoded gene transcripts, were not affected in our mutant RNA-seq datasets, the observed OXPHOS deficiency phenotype may progress directly through mito-translation but not through mito-transcription. Some aspects of the cardiac phenotype such as LVNC are similar to the published Hif1a_90_, Vhl_66_, or Prdm16_85_ mutants; however, the reported transcriptomes are dissimilar to our Pcbp1-cKO or Aars2-cKO, and do not elicit a ISR/UPR signature as do our mutants. Our result indicates a uniform UPR/ISR signature in both of our mutants with up-regulation of Atf4, Ddit3, Asns, as wells as a concordant up-regulation of cytosolic ARS genes. The shift in expression of these specific genes suggests a potential transcription regulatory mechanism downstream of the the PCBP1-AARS2 axis. The majority of the genes encoding cytosolic ARS contains C/ebp-Atf-Response-Element (CARE) enhancers and the induction of the cytosolic ARS under amino acid limitation stress is thought to be induced by Atf4_65_. Asns, a UPR/ISR gene potently up-regulated in both of the Pcbp1 and Aars2 mutant hearts also contain the CARE enhancer and is a target of Atf4_54_. Together, this would suggest the PCBP1-AARS2 axis induced UPR/ISR potentially through the dysregulation in mito-translation, and activates the nuclear transcription factor Atf4. There are similarities for this mito-nuclear retrograde signaling in the heart as demonstrated by a recent report where mitochondrial translation defects lead to a repression of nuclear transcription program, through metabolites L-phenylalanie and AMP_91_. Together, our finding indicates a unique functional specificity regulated by the PCBP1-AARS2 axis in cardiomyocyte development and contribute to our further understanding of mitochondrial cardiomyopathy.

In conclusion, our finding provides new insight into the mechanism whereby the RBP PCBP1 regulates the alternative splicing of a nuclear encoded mitochondrial tRNA aminoacyl transferase AARS2, a gene previously identified to be causal for a form of mitochondrial cardiomyopathy (COXPD8, OMIM: 614096). Our results provide clear evidence that this newly identified PCBP1-AARS2 axis safeguards mitochondrial translation and stabilizes OXPHOS complexes in the heart. There is currently no efficient treatment for OXPHOS deficiency diseases, and gaining new knowledge into these tissue-specific regulatory mechanisms may reveal potential targets for therapeutic intervention. Further research is needed to determine the translational targets and metabolic pathways governed by this novel axis in heart development and congenital heart diseases.

## RESOURCE AVAILABILITY

### Lead contact

Further information and reasonable requests for resources and reagents should be directed to the Lead Contact, Da-Zhi Wang

### Materials availability

Materials used in this study are available from the Lead Contact upon reasonable request. RNA-seq and eCLIP-seq data have been deposited at GEO and are publicly available as of the date of publication. Accession numbers are listed in the key resources table. Mouse lines generated in this study are available from the Lead Contact upon reasonable request.

## EXPERIMENTAL MODELS AND SUBJECT DETAILS

### Animal husbandry and gene manipulations

CRISPR/Cas9 pronuclear microinjection was used to generate Aars2-Flox and deletion mutant mice at the Mouse Gene Manipulation Core of Boston Children’s Hospital as described_92_ with minor modifications. Briefly, a cocktail of 0.61pmol/ul each of crRNA+tracrRNA, 100ng/ul Cas9 protein and 10ng/ul ssODN (IDT) or ds-Donor DNA was injected into the pronuclei of E0.5 embryos (C57Bl6/Harlan). Post-injection survived embryos were re-implanted into recipient CD1 pseudo-pregnant females and allowed to develop to term. The resultant pups were genotyped by PCR, and genome-edited founders identified and verified by TOPO-cloning followed by Sanger sequencing. Two founders were chosen from each line and backcrossed to C57BL/6 (Harlan) for three generations to establish lines. Genotyping primers for screening founders are listed in **Supplemental Table 6**.

We obtained a Pcbp1-Flox allele_17_ for conditional knockout generation. Mice carrying Pcbp1-Flox or Aars2-Flox were bred with Tg(Tnnt2-cre)5Blh/JiaoJ (Ctnt-Cre)_34_ to generate corresponding cardiac-specific conditional deletions (Pcbp1-cKO or Aars2-cKO). All data presented here for Pcbp1-cKO are from the previously described non-hypomorphic Pcbp1-Flox allele_17_, although we had also examined our independently generated hypomorphic Pcbp1-Flox allele_21_ for conditional knockout, and observed a very similar phenotype.

All animals were maintained in a 12-hour light cycle. Genetically modified strains of mice were genotyped by isolating genomic DNA from a tail tip excision and subjecting the DNA to PCR reaction using primers corresponding to the modification. For timed pregnancy experiments, two months old male and female from indicated lines and genotypes were be co-housed, and vaginal plugs of the female mice checked each morning following the date of co-housing to approximate the date of gestation, with the day of vaginal plug appearance noted as E0.5. Mice were euthanized by CO2 asphyxiation according to approved protocols.

All murine experiments were performed in accordance with approved protocols of the Boston Children’s Hospital Institutional Animal Care and Use Committee.

## METHOD DETAILS

### High-resolution Episcopic Microscopy and 3D reconstruction

Mouse embryos were processed for high-resolution episcopic microscopy as described previously_37_. Briefly, E12.5 embryos were fixed in 10% neutral buffered formalin solution at 4°C for 72 hours. Embryos were then incubated in clearing solution (4M urea, 10% Glycerol, 4% Sodium Dodecyl Sulfate in double distilled water) for 8 days in room temperature, and the solution replaced every 2 days. Cleared embryos were subsequently washed in PBS for 3 times for 5 minutes each, and dehydrated in ascending ethanol series (50%, 60%, 70%, 80%, 85%, 90%, 95%, and 100% ethanol) at room temperature with gentle rocking, for 2 hours at each step. The dehydrated embryos were stained with staining solution (0.1375 g Eosin Y Disodium, 0.0275 g Acridine Orange hemi(zinc chloride) salt in 50ml of ethanol) for 1 hour, and then replaced with fresh staining solution to stain overnight. Infiltration solution (100 ml Solution A of JB-4 embedding kit, 1.25 g Catalyst C, 0.275 g Eosin Y Disodium Salt, 0.055 g Acridine Orange hemi(zinc chloride) salt) was made, and the stained embryos were first incubated in a 1:1 mixture of staining solution and infiltration solution for 3 hours in 4°C, then in 100% infiltration solution for 3 hours in 4°C, then in fresh infiltration solution for 72 hours in 4°C. The solution was changed every 24 hours. The processed embryos were then embedded in resin material as described previously_37_, and the embedded blocks were mounted on a microtome. The cut face was aligned with a stereo zoom microscope horizontally with the camera acquisition based on excitation 470 ± 20 nm, dichroic 495 nm, emission 525 ± 25 nm. Each section was cut at 2.5 μm, and with each cut an image was acquired, until the entire embryo was completed. 3D reconstruction, visualization, and measurement were performed using Amira software (2019.1, ThermoFisher).

### Histology

Mouse heart tissues were dissected out, rinsed with PBS, and fixed in 4% paraformaldehyde (pH 7.4) overnight. After dehydration through a series of ethanol baths, samples were embedded in paraffin wax according to standard laboratory procedures. Sections of 5 μm were stained with haematoxylin and eosin (H&E). The stained sections were used for routine histological examination by light microscopy and quantified with ImageJ software.

### Immunofluorescence labeling

Mouse embryos and embryonic hearts were harvested at different stages and fixed in ice cold PBS containing 4% paraformaldehyde overnight at 4°C. Fixed embryos were washed with PBS, then saturated in a 30% sucrose solution in PBS for 6 hours in 4°C, followed by a 30 minutes incubation with 1:1 OCT(Tissue-Tek):30% Sucrose, and embedded in OCT. Frozen blocks were then cut into 8-μm sections using a Leica CM 3050S cryostat (Leica Microsystems).

For paraffin embedded tissue section, they were first de-paraffined and rehydrated using xylene follow by wash in descending ethanol series (100%, 95%, 70%, 50%) and water. De-paraffined slides were then processed for antigen retrieval using Retrievagen A (pH 6.0) (BD Bioscience) according to manufacturer’s protocol. Antigen-retrieved slides were then processed for immunofluorescence staining as below.

Sections were permeabilized with wash buffer (PBS with 0.2% Triton 100), then blocked with blocking buffer (PBS with 1% BSA, 5% donkey serum, 0.2%Triton 100) for 1h at room temperature, followed by overnight incubation at 4°C with primary antibody diluted in blocking buffer. Sections were rinsed in wash buffer and incubated with fluorescently conjugated secondary antibodies and DAPI diluted in blocking buffer at room temperature for 1 hour, before rinsing and mounting in the Prolong Glass mounting medium (Thermo Fisher). Sections were imaged using an Olympus Fluoview FV3000 confocal microscope. n=4 hearts per group, and at least 4 sections of the heart were imaged and examined. Detailed antibody information is provided in the **Supplemtal Table 7**.

### mRNA *in situ* hybridization

Single mRNA molecule in situ hybridization coupled with immunofluorescence staining were performed according to the manufacture protocol of the RNAscope® 2.5 HD (RED) Assay (Advanced Cell Diagnostics, cat# 322360), or RNAscope® Multiplex Fluorescent Reagent Kit v2 (Advanced Cell Diagnostics, cat# 323100) with modifications by a previously published protocol_40_. Briefly, embryonic heart tissues were fixed in 4% paraformaldehyde in 4°C for 24 hours, and were either embedded in OCT compound or paraffin (antigen retrieved) as described in the previous section. RNA hybridization was performed first according to manufacturer’s protocol, followed by immunofluorescence staining. The following RNAscope probes were used: Mm-Hey2 (cat# 404651), Mm-Nppa (cat# 418691), Mm-Myl4 (cat# 443801), Mm-Myl7-C3 (cat# 584271-C3), Mm-Bmp10 (cat# 415921). All *in situ* hybridization was performed on n=3 hearts per group, and at least 4 sections of the heart were imaged and examined with an Olympus Fluoview FV3000 confocal microscope.

### Western blotting

Embryonic heart ventricles at E16.5 were isolated, and protein was extracted using SDS lysis buffer_93_ with cOmplete protease inhibitor cocktail (Roche) and PhosSTOP phosphatase inhibitor cocktail (Roche), with mortar and pestle on ice for complete tissue lysis. The total protein was measured by BCA protein assay (Pierce), and the protein loading was normalized accordingly. Diluted protein lysates in Laemmli buffer were incubated at 95°C for 10 minutes, and were subsequently separate by SDS-PAGE and transferred to 0.45 μm nitrocellulose membranes. The membranes were blocked with 5% BSA in PBS with Tween-20 (PBST) for 1 hour in room temperature, and then incubated with primary antibodies at 4°C for 16 hours. The membranes were then washed 3x10 minutes in PBST, and incubated with HRP-conjugated secondary antibodies accordingly for 1.5 hours in room temperature. Membranes were then washed for 4x10 minutes in PBST. The immunoreactive proteins were detected by enhanced chemiluminescence with autoradiography. Western blotting densitometry quantifications were performed with NIH ImageJ software with n=3. Detailed antibody information is provided in the **Supplemental Table 7**.

### Echocardiography

Echocardiographic measurements were performed on neonatal mice using a Visualsonics Vevo® 2100 Imaging System (Visual Sonics, Toronto, Canada) with a 40 MHz MicroScan transducer (model MS-550D). Heart rate and LV dimensions, including diastolic and systolic wall thicknesses, LV end-diastolic and end-systolic chamber dimensions were measured from 2-D short-axis under M-mode tracings at the level of the papillary muscle. LV mass and functional parameters such as percentage of fractional shortening (FS%) and ejection fraction (EF%) were calculated using the above primary measurements and accompanying software.

### Reverse transcription and quantitative PCR analysis

Total RNAs were isolated using TRIzol Reagent (Life Technologies) from cells and tissue samples. For quantitative RT-PCR, 2 μg RNA samples were reverse-transcribed to cDNA using random hexamers and MMLV reverse transcriptase (Life Technologies) in 20 μl reactions. In each analysis, 0.1 μl cDNA pool was used for quantitative PCR. Real-time PCR was performed using an ABI 7500 thermocycler with Power SYBR Green PCR Kit (Thermo Fisher, 4368702). The relative expression of interested genes is normalized to the expression of 18S rRNA or β-actin. qRT-PCR primers are listed in **Supplemental Table 5**.

### RNA-sequencing and data analysis

Total RNA from the ventricles of embryonic day (E) 16.5 hearts were isolated using TRIzol (Life Technologies) as described, and strand-specific polyA-selected libraries were generated with TruSeq RNA Library Preparation Kit (Illumina). The obtained libraries were combined in an equimolar amount and loaded on Illumina flow cell lanes, followed by paired-end sequencing (PE150bp, GENEWIZ), to achieve at least 80 million reads per sample. FASTQ files were extracted, and the TruSeq sequencing adapters and low-quality reads were removed from FASTQ files with Cutadapt.

For gene level analysis, the cleaned FASTQ files were quality checked using FastQC (Babraham Bioinformatics), then aligned to mouse genome (Esembl GRCm38 genome obtained from GENCODE) using HISAT2 (v.2.1.0)_94,95_. Subsequently, transcript assembly was performed using StringTie (v.1.3.4)_96_ with annotated transcriptome as reference. The assembled transcriptomes were quantified using prepDE.py script provided by the StringTie developer to generate gene matrix files. EdgeR (v.3.26.1)_97_ was used to compute Counts Per Million (CPM) as a normalized measurement for gene expression. Differentially expressed genes were tested using Fisher’s exact test, and multiplicity correction was performed with the Benjamini-Hochberg method on the p-values to control the false discovery rate (FDR). Differentially regulated genes with FDR < 0.05 were considered significant.

For alternative splicing analysis, the cleaned FASTQ files were quality checked using FastQC (Babraham Bioinformatics), then aligned to the mouse genome (Esembl GRCm38 genome obtained from GENCODE) using STAR (2.7.3) with the following setting: “--outFilterType BySJout --outFilterMultimapNmax 20 --alignSJoverhangMin 8 -- alignSJDBoverhangMin 1 --outFilterMismatchNmax 999 --alignIntronMin 10 -- alignIntronMax 1000000 --alignMatesGapMax 1000000”. The aligned splice junctions detected from the first run from each sample were subsequently inserted to the STAR alignment run the second time to detect novel splice junctions. Splice graphs for local splice variation were generated with MAJIQ (2.1)_98_. Alternative splicing related motifs enrichment was performed with rMATs (4.0.2)_58_ follow by rMAPs (2.0.0)_61_ with default parameters.

### Principal Component Analysis and Gene Set Enrichment Analysis

We defined expressed genes as those that have expression in at least half of all samples. Expressed genes were subjected to Principal Component Analysis (PCA). Principal component 1 and 2 were plotted in 2-D coordinates. Gene Set Enrichment Analysis (GSEA) were performed on expressed genes according to software manual_99,100_. Gene sets with nominal p-value < 0.05 and FDR < 0.25 were considered significant. All expressed genes were Log2 transformed, centered, and unsupervised hierarchical clustering were performed using k-mean clustering method with Cluster 3.0 software_101_. Java Treeview (v.3.0) was used to visualize the clustered heatmaps.

### Enhance Crosslinking and Immunoprecipitation follow by sequencing (eCLIP-seq)

eCLIP studies were performed according to the published single-end seCLIP protocol_102_ with the following modifications (Eclipse Bioinnovations Inc). Approximately 80 mg of frozen embryonic mouse heart tissue was cryogrinded and UV crosslinked at 400 mJoules/cm2 with 254 nm radiation. The equivalence of 75 μg of RNA was lysed using 1 mL of eCLIP lysis mix and subjected to two rounds of sonication for 4 minutes with 30 second ON/OFF at 75% amplitude. A pre-validated PCBP1 antibody (Abcam cat# ab168377) was then pre-coupled to Anti-Rabbit IgG Dynabeads (ThermoFisher), added to lysate, and incubated overnight at 4°C. Prior to immunoprecipitation, 2% of the sample was taken as the paired input sample, with the remainder magnetically separated and washed with eCLIP high stringency wash buffers. IP and input samples were cut from the membrane at the relative band size to 75kDa above. RNA adapter ligation, IP-western, reverse transcription, DNA adapter ligation, and PCR amplification were performed as previously described_28_.

The eCLIP cDNA adapter contains a sequence of 10 random nucleotides at the 5’ end. This random sequence serves as a unique molecular identifier (UMI)_103_ after sequencing primers are ligated to the 3’ end of cDNA molecules. Therefore, eCLIP reads begin with the UMI and, in the first step of analysis, UMIs were pruned from read sequences using umi_tools (v0.5.1)_104_. UMI sequences were saved by incorporating them into the read names in the FASTQ files to be utilized in subsequent analysis steps. Next, 3’-adapters were trimmed from reads using cutadapt (v2.7)_105_, and reads shorter than 18 bp in length were removed. Reads were then mapped to a database of human and mouse repetitive elements and rRNA sequences compiled from Dfam_106_ and Genbank_107_. All non-repeat mapped reads were mapped to the mouse genome (mm10) using STAR (v2.6.0c)_108_. PCR duplicates were removed using umi_tools (v0.5.1) by utilizing UMI sequences from the read names and mapping positions. Peaks were identified within eCLIP samples using the peak caller CLIPper (https://github.com/YeoLab/clipper)^29^. For each peak, IP versus input fold enrichment values were calculated as a ratio of counts of reads overlapping the peak region in the IP and the input samples (read counts in each sample were normalized against the total number of reads in the sample after PCR duplicate removal). A p-value was calculated for each peak by the Yates’ Chi-Square test, or Fisher Exact Test if the observed or expected read number was below 5. Comparison of different sample conditions was evaluated in the same manner as IP versus input enrichment; for each peak called in IP libraries of one sample type we calculated enrichment and p-values relative to normalized counts of reads overlapping these peaks in another sample type. Peaks were annotated using transcript information from GENCODE_109_ with the following priority hierarchy to define the final annotation of overlapping features: protein coding transcript (CDS, UTRs, intron), followed by non-coding transcripts (exon, intron). De novel motif discovery for peaks detected were performed using Homer (v4.11), and the positionally enriched k-mer analysis (PEKA) in whole gene, 3’UTR, intron or exon regions was performed using PEKA software (https://github.com/ulelab/peka)^33^.

### RNA-Binding Protein Immunoprecipitation (RIP) followed by qRT-PCR

RNA-Binding protein immunoprecipitation (RIP) for PCBP1 was perform using Magna RIP RNA-binding protein immunoprecipitation kit (Millipore, cat 17-700) and the same PCBP1 antibody used for eCLIP-seq (Abcam cat# ab168377), according to manufacturer’s protocol. Briefly, heart tissues were snap frozen in liquid nitrogen, and quickly ground into powder within the liquid nitrogen bath. Ground tissues were lysed in RIP Lysis Buffer (Millipore, cat# CS203176) on ice. 10 μl of the supernatant of the RIP lysate was collected as input control. Magnetic Bead and antibody (Pcbp1 antibody or rabbit IgG) were incubated for 1 hour, before adding the antibody bound magnetic beads into the lysate for immunoprecipitation. The RBP immunoprecipitation mixture was incubated at 4°C overnight. RIP lysate was separated by a magnetic separator, and the beads washed with ice-cold RIP wash buffer (Millipore cat# CS203177) for 6 times. The washed RIP complexes were treated with Proteinase K for 30 minutes at 55°C, washed once with RIP wash buffer, and proceeded to pheno:chloroform:isoamyl alcohol extraction of bound RNA. RNA from the input sample was processed at the same time. The quantity and quality of the RNA isolated from Pcbp1-RIP, IgG-RIP, and input were assessed by NanoDrop, and processed for qRT-PCR. RIP RNA fractions were normalized to input RNA fraction by: ΔCt [normalized RIP] = (Ct [RIP] – (Ct [Input] – Log2 (Input Dilution Factor)), and the fold enrichment for RIP-fraction is calculated as % Input = 2^(–ΔCt [normalized RIP] – ΔCt [normalized IgG])^. 3 neonatal P0 heart ventricles from wild-type pups were pulled into one sample, and the experiment was performed in triplicate (n=3, 9 hearts total).

### Cell Culture

The cardiac muscle cell line HL-1 was cultured and maintained as described^110^. Briefly, HL-1 cells were cultured in Claycomb medium (Sigma Aldrich) supplemented with 10% Fetal Bovine Serum (FBS) (Sigma Aldrich), 0.1mM Norepinephrine (Sigma Aldrich), 2mM L-Glutamine and Penicillin/Streptomycin (100U/ml and 100ug/ml). Supplemented Claycomb media were changed every day, and cells were split before they reached confluence.

### Statistics

Values are reported as means ± SEM unless indicated otherwise. One-way analysis of variance analysis (ANOVA) followed by Turkey Post-Hoc testing was used to evaluate the statistical significance for multiple-group comparisons. In addition, the 2-tailed non-parametric Student’s T-test was used for 2-group comparisons. Values of P<0.05 were considered statistically significant.

### Data and code availability

All data that support the findings of this study are available within the article and its supplemental information files, and from the corresponding authors upon reasonable request.

## Acknowledgements

We thank members of the Da-Zhi Wang Lab and the Hong Chen Lab for their help and scientific discussion. We appreciate the insightful feedback provided by Drs. William Pu and Jonathan Seidman and Yuxuan Guo and Yangpo Cao in the Pu lab for sharing antibody reagents. We thank the imaging core facilities at Boston Children’s Hospital and the administrative staff in the Department of Cardiology and the Vascular Biology Program for their support. We also would like to thank the IDDRC Gene Manipulation Core at the Boston Children’s Hospital for their help with performing microinjections for generation of the Aars2-Flox mouse line. This research was conducted with assistance from the HMS O2 High Performance Compute Cluster, supported by the Research Computing Group at Harvard Medical School.

