## Supplemental Figures for "PCBP1 regulates alternative splicing of AARS2 in congenital cardiomyopathy"

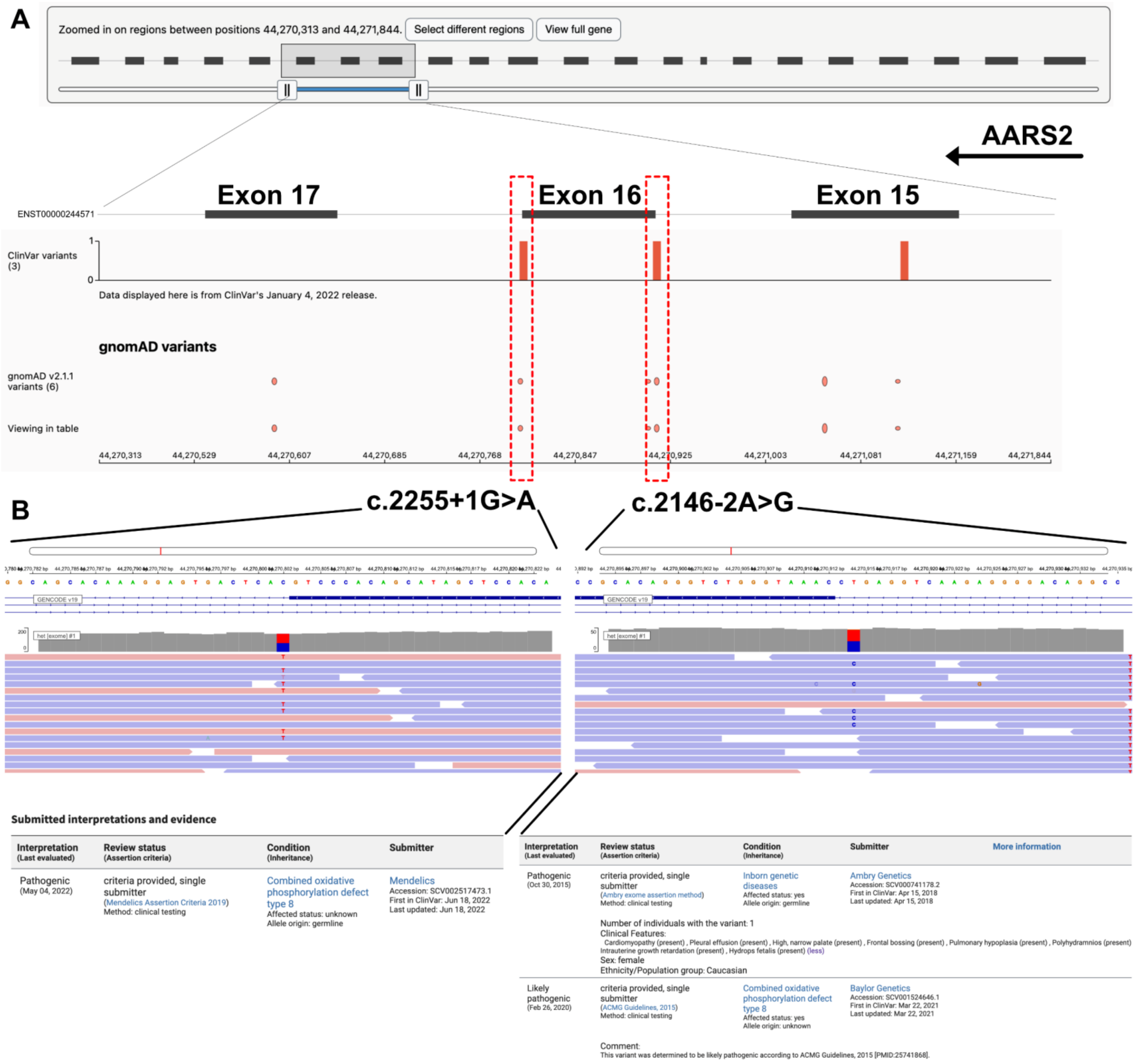
A
Zoomed in on regions between positions 44,270,313 and 44,271,844.
Select different regions
View full gene
AARS2
Exon 17
Exon 16
Exon 15
ENST00000244571
ClinVar variants (3)
1
0
Data displayed here is from ClinVar's January 4, 2022 release.
gnomAD variants
gnomAD v2.1.1 variants (6)
Viewing in table
44,270,313
44,270,529
44,270,607
44,270,685
44,270,768
44,270,847
44,270,925
44,271,003
44,271,081
44,271,159
44,271,844
B
c.2255+1G>A
c.2146-2A>G
GENCODE v19
het [exome] #1
Submitted interpretations and evidence
Interpretation (Last evaluated)
Review status (Assertion criteria)
Condition (Inheritance)
Submitter
Pathogenic (May 04, 2022)
criteria provided, single submitter (Mendelics Assertion Criteria 2019) Method: clinical testing
Combined oxidative phosphorylation defect type 8
Affected status: unknown
Allele origin: germline
Mendelics
Accession: SCV002517473.1
First in ClinVar: Jun 18, 2022
Last updated: Jun 18, 2022
More information
Pathogenic (Oct 30, 2015)
criteria provided, single submitter (Ambry exome assertion method) Method: clinical testing
Inborn genetic diseases
Affected status: yes
Allele origin: germline
Ambry Genetics
Accession: SCV000741178.2
First in ClinVar: Apr 15, 2018
Last updated: Apr 15, 2018
Number of individuals with the variant: 1
Clinical Features:
Cardiomyopathy (present) , Pleural effusion (present) , High, narrow palate (present) , Frontal bossing (present) , Pulmonary hypoplasia (present) , Polyhydramnios (present) Intrauterine growth retardation (present) , Hydrops fetalis (present) (less)
Sex: female
Ethnicity/Population group: Caucasian
Likely pathogenic (Feb 26, 2020)
criteria provided, single submitter (ACMG Guidelines, 2015) Method: clinical testing
Combined oxidative phosphorylation defect type 8
Affected status: yes
Allele origin: unknown
Baylor Genetics
Accession: SCV001524646.1
First in ClinVar: Mar 22, 2021
Last updated: Mar 22, 2021
Comment:
This variant was determined to be likely pathogenic according to ACMG Guidelines, 2015 [PMID:25741868].

**Supplemental Figure 1. Pathogenic variants located at splice sites flanking exon 16 of AARS2.** (A)Two pathogenic variants located at the intronic splice donor (c.2255+1G>A) and acceptor site (c.2146-2A>G) were shown. (B) The details of these variants were shown within the NCBI genome browser, including the descriptions from the NCBI ClinVar database.

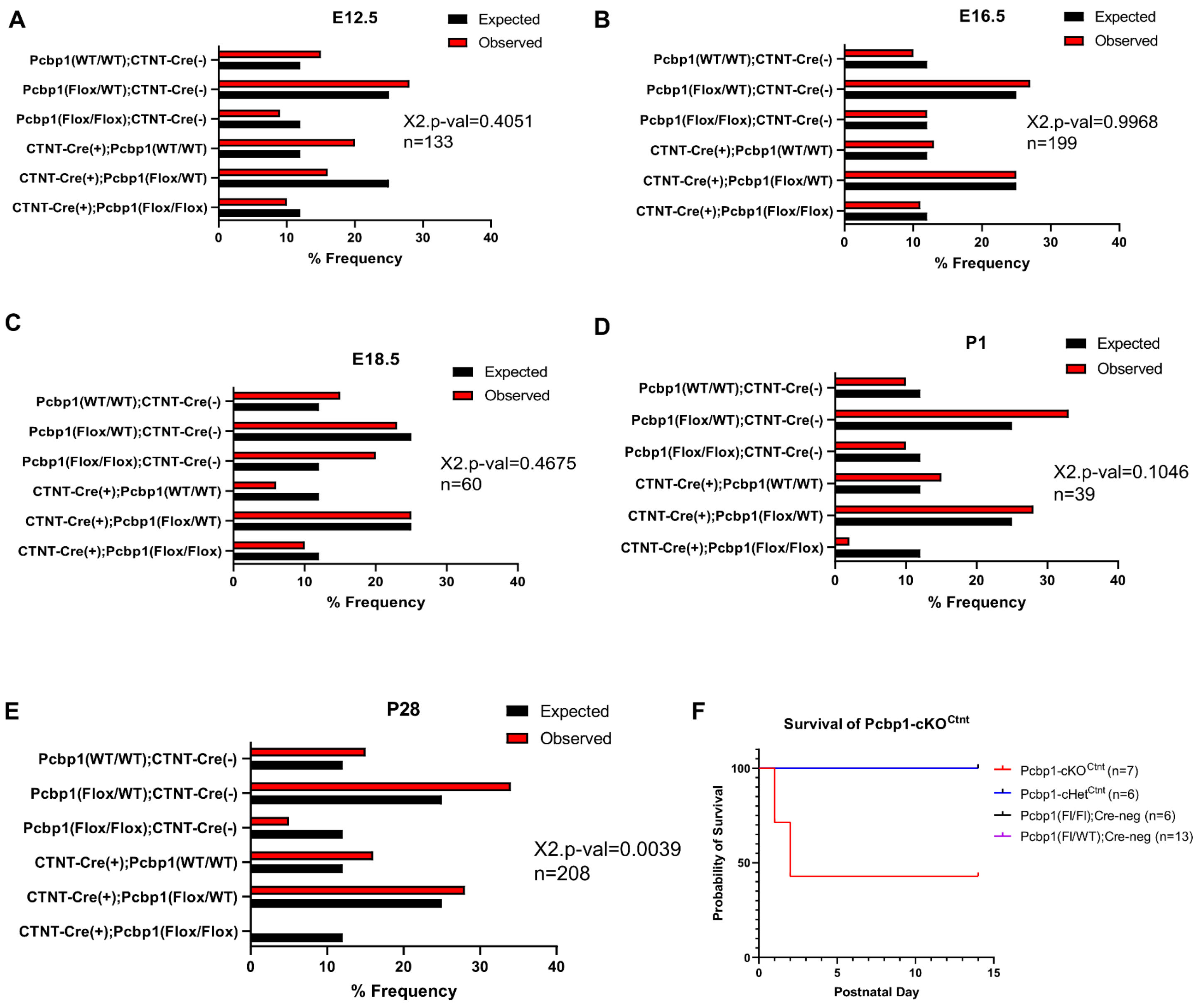
A
E12.5
Expected
Observed
Pcbp1(WT/WT);CTNT-Cre(-)
Pcbp1(Flox/WT);CTNT-Cre(-)
Pcbp1(Flox/Flox);CTNT-Cre(-)
CTNT-Cre(+);Pcbp1(WT/WT)
CTNT-Cre(+);Pcbp1(Flox/WT)
CTNT-Cre(+);Pcbp1(Flox/Flox)
X2.p-val=0.4051
n=133
0
10
20
30
40
% Frequency
B
E16.5
Expected
Observed
Pcbp1(WT/WT);CTNT-Cre(-)
Pcbp1(Flox/WT);CTNT-Cre(-)
Pcbp1(Flox/Flox);CTNT-Cre(-)
CTNT-Cre(+);Pcbp1(WT/WT)
CTNT-Cre(+);Pcbp1(Flox/WT)
CTNT-Cre(+);Pcbp1(Flox/Flox)
X2.p-val=0.9968
n=199
0
10
20
30
40
% Frequency
C
E18.5
Expected
Observed
Pcbp1(WT/WT);CTNT-Cre(-)
Pcbp1(Flox/WT);CTNT-Cre(-)
Pcbp1(Flox/Flox);CTNT-Cre(-)
CTNT-Cre(+);Pcbp1(WT/WT)
CTNT-Cre(+);Pcbp1(Flox/WT)
CTNT-Cre(+);Pcbp1(Flox/Flox)
X2.p-val=0.4675
n=60
0
10
20
30
40
% Frequency
D
P1
Expected
Observed
Pcbp1(WT/WT);CTNT-Cre(-)
Pcbp1(Flox/WT);CTNT-Cre(-)
Pcbp1(Flox/Flox);CTNT-Cre(-)
CTNT-Cre(+);Pcbp1(WT/WT)
CTNT-Cre(+);Pcbp1(Flox/WT)
CTNT-Cre(+);Pcbp1(Flox/Flox)
X2.p-val=0.1046
n=39
0
10
20
30
40
% Frequency
E
P28
Expected
Observed
Pcbp1(WT/WT);CTNT-Cre(-)
Pcbp1(Flox/WT);CTNT-Cre(-)
Pcbp1(Flox/Flox);CTNT-Cre(-)
CTNT-Cre(+);Pcbp1(WT/WT)
CTNT-Cre(+);Pcbp1(Flox/WT)
CTNT-Cre(+);Pcbp1(Flox/Flox)
X2.p-val=0.0039
n=208
0
10
20
30
40
% Frequency
F
Survival of Pcbp1-cKO^Ctnt
Probability of Survival
100
50
0
0
5
10
15
Postnatal Day
Pcbp1-cKO^Ctnt (n=7)
Pcbp1-cHet^Ctnt (n=6)
Pcbp1(Fl/Fl);Cre-neg (n=6)
Pcbp1(Fl/WT);Cre-neg (n=13)

**Supplemental Figure 2. Observed frequency for Pcbp1-cKO mice compared to littermate controls at different developmental stages.** Myocardial deletion of Pcbp1 was generated by crossing cTNT-Cre(+); Pcbp1(Flox/WT) with Pcbp1(Flox/WT) mice, 6 possible genotypes were generated as indicated in (A-E). Observed and expected frequency for E12.5 (A), E16.5 (B), E18.5 (C), P1 (D), and P28 (E) were shown as indicated, and Chi-square test was performed to examine the difference between the observed and expected results at each developmental stage. (F) To examine postnatal survival, myocardial deletion of Pcbp1 was generated by crossing cTNT-Cre(+); Pcbp1(Flox/WT) with Pcbp1(Flox/Flox) mice, and 4 possible genotypes were generated as indicated in (F), where a Kaplan-Meier estimate of probability of survival was plotted for these genotypes within two weeks postnatally.

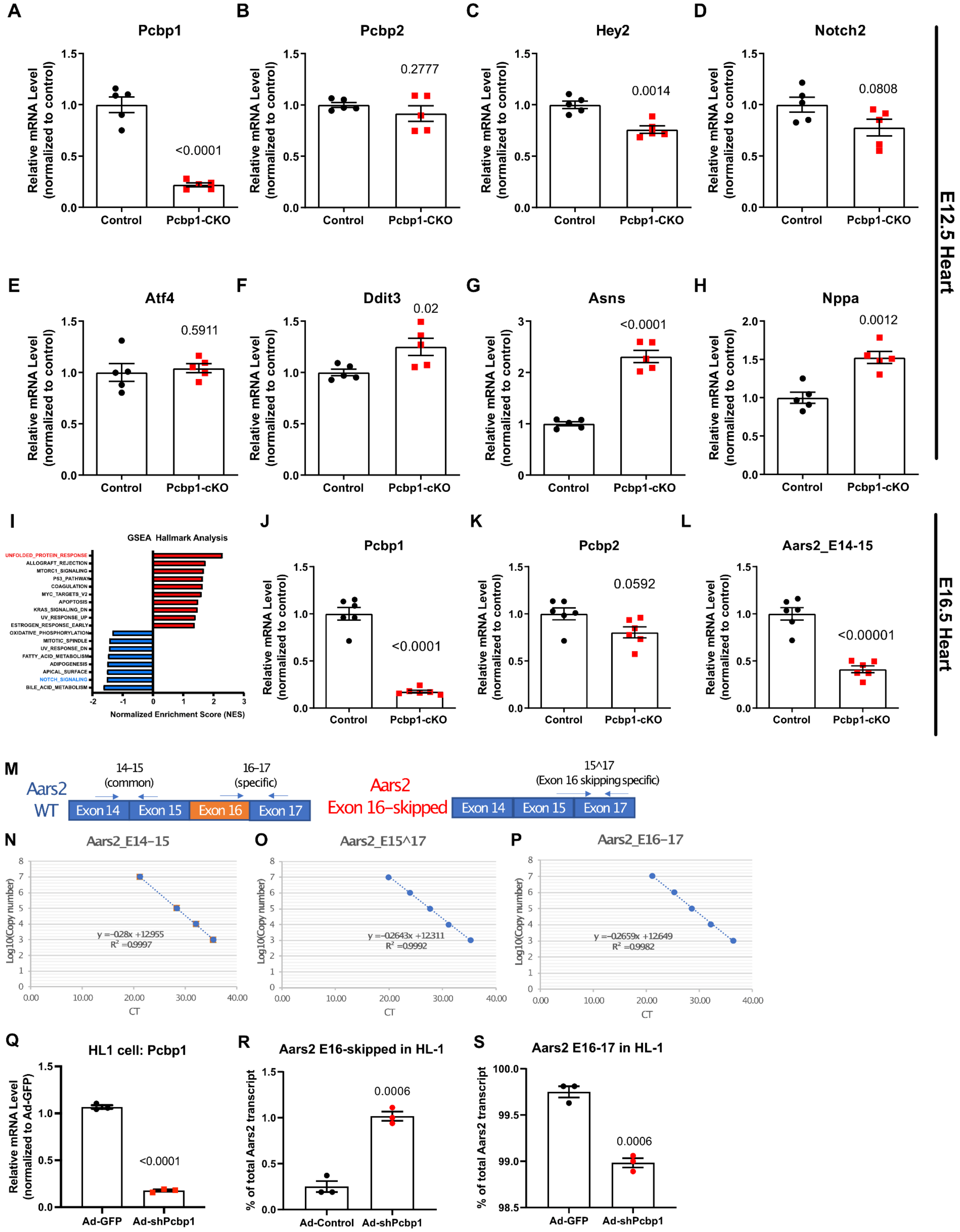
A
Pcbp1
Relative mRNA Level (normalized to control)
<0.0001
Control
Pcbp1-CKO
B
Pcbp2
0.2777
C
Hey2
0.0014
D
Notch2
0.0808
E12.5 Heart
E
Atf4
0.5911
Pcbp1-cKO
F
Ddit3
0.02
G
Asns
<0.0001
H
Nppa
0.0012
I
GSEA Hallmark Analysis
UNFOLDED_PROTEIN_RESPONSE
ALLOGRAFT_REJECTION
MTORC1_SIGNALING
P53_PATHWAY
COAGULATION
MYC_TARGETS_V2
APOPTOSIS
KRAS_SIGNALING_DN
UV_RESPONSE_UP
ESTROGEN_RESPONSE_EARLY
OXIDATIVE_PHOSPHORYLATION
MITOTIC_SPINDLE
UV_RESPONSE_DN
FATTY_ACID_METABOLISM
ADIPOGENESIS
APICAL_SURFACE
NOTCH_SIGNALING
BILE_ACID_METABOLISM
Normalized Enrichment Score (NES)
J
Pcbp1
<0.0001
K
Pcbp2
0.0592
L
Aars2_E14-15
<0.00001
E16.5 Heart
M
Aars2 WT
14–15 (common)
16–17 (specific)
Exon 14
Exon 15
Exon 16
Exon 17
Aars2 Exon 16–skipped
15^17 (Exon 16 skipping specific)
N
Aars2_E14–15
Log10(Copy number)
CT
y = -0.28x + 12.955
R² = 0.9997
O
Aars2_E15^17
y = -0.2643x + 12.311
R² = 0.9992
P
Aars2_E16–17
y = -0.2659x + 12.649
R² = 0.9982
Q
HL1 cell: Pcbp1
Relative mRNA Level (normalized to Ad-GFP)
<0.0001
Ad-GFP
Ad-shPcbp1
R
Aars2 E16-skipped in HL-1
% of total Aars2 transcript
0.0006
Ad-Control
Ad-shPcbp1
S
Aars2 E16-17 in HL-1
% of total Aars2 transcript
0.0006
Ad-GFP
Ad-shPcbp1

**Supplemental Figure 3. Gene expression measurements for E12.5 and E16.5 Pcbp1-cKO hearts.** qRT-PCR was performed on E12.5 RNA from Pcbp1-cKO and control hearts. Pcbp1 (A), Pcbp2 (B), Hey2 (C), Notch2 (D), Atf4 (E), Ddit3 (F), Asns (G), and Nppa (H) were assayed (n=5, Student's T-Test p-values are shown). (I) GSEA Hallmark analysis result for E16.5 Pcbp1-cKO vs Control hearts were shown. qRT-PCR was performed on E16.5 RNA from Pcbp1-cKO and control hearts, and Pcbp1 (J), Pcbp2 (K), and the exon14-15 region of Aars2 (L) were shown. (n=6, Student's T-Test p-values are shown). (M) Absolute qRT-PCR with corresponding amplicons were developed as indicated to quantify the common exon junction (E14-15) as an estimate for total transcript, as well as primers specifically detecting Exon 16-17 and Exon 15^17 (E16 skipped) transcripts. (N) Absolute qRT-PCR with corresponding amplicons with serial dilutions of known copy numbers were performed to generate standard curve equations as shown in (N-P), which were then used to estimate the absolute copy number for each of these exon-junctions present in the samples. (Q) Knockdown of Pcbp1 by adenoviral expression of shRNA targeting Pcbp1 was performed in HL-1 cells, and RNA was isolated 72 hours post-infection to assess Pcbp1 knockdown levels by qRT-PCR. Aars2 exon 16 skipping was assessed by absolute qRT-PCR by E15^17 specific primers (R) and E16-17 specific primers (S) (n=3, Student's T-Test p-values are shown).

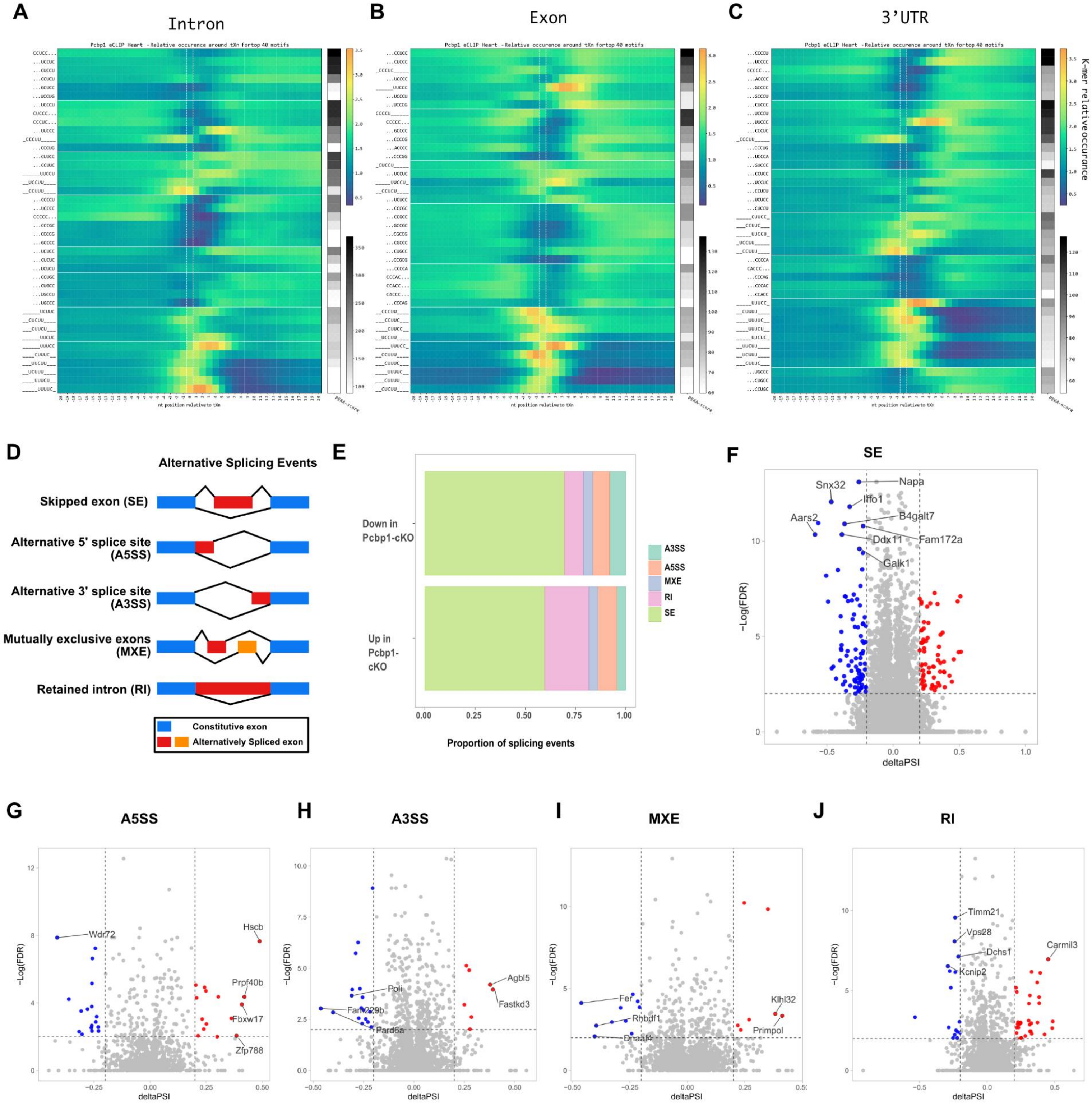

A
Intron
B
Exon
C
3'UTR
Pcbp1 eCLIP Heart - Relative occurence around tXn fortop 40 motifs
nt position relative to tXn
PEKA-score
K-mer relativeoccurance
D
Alternative Splicing Events
Skipped exon (SE)
Alternative 5' splice site (A5SS)
Alternative 3' splice site (A3SS)
Mutually exclusive exons (MXE)
Retained intron (RI)
Constitutive exon
Alternatively Spliced exon
E
Down in Pcbp1-cKO
Up in Pcbp1-cKO
A3SS
A5SS
MXE
RI
SE
0.00
0.25
0.50
0.75
1.00
Proportion of splicing events
F
SE
Snx32
Napa
Iffo1
Aars2
B4galt7
Ddx11
Fam172a
Galk1
−Log(FDR)
deltaPSI
G
A5SS
Wdr72
Hscb
Prpf40b
Fbxw17
Zfp788
H
A3SS
Agbl5
Fastkd3
Poli
Fam229b
Pard6a
I
MXE
Fer
Rhbdf1
Dnaaf4
Klhl32
Primpol
J
RI
Timm21
Vps28
Dchs1
Carmil3
Kcnip2

**Supplemental Figure 4. Detailed results for the positionally enriched k-mer analysis on PCBP1 eCLIP-seq, and alternative splicing events detected in Pcbp1-cKO heart.** Positionally enriched k-mer analysis was performed on PCBP1 eCLIP-seq peaks localized on intron (A), exon (B), or 3'UTR (C) regions. (D) Alternative splicing events from the E16.5 RNA-seq data with Pcbp1-cKO and control were detected by rMATS, corresponding to five major types of alternative splicing patterns as indicated. (E) Summary of the proportion of the differential splicing events from each major alternative splicing pattern. (F-J) Volcano plots for the delta PSI (percent spliced in) were shown for each major splicing patterns.

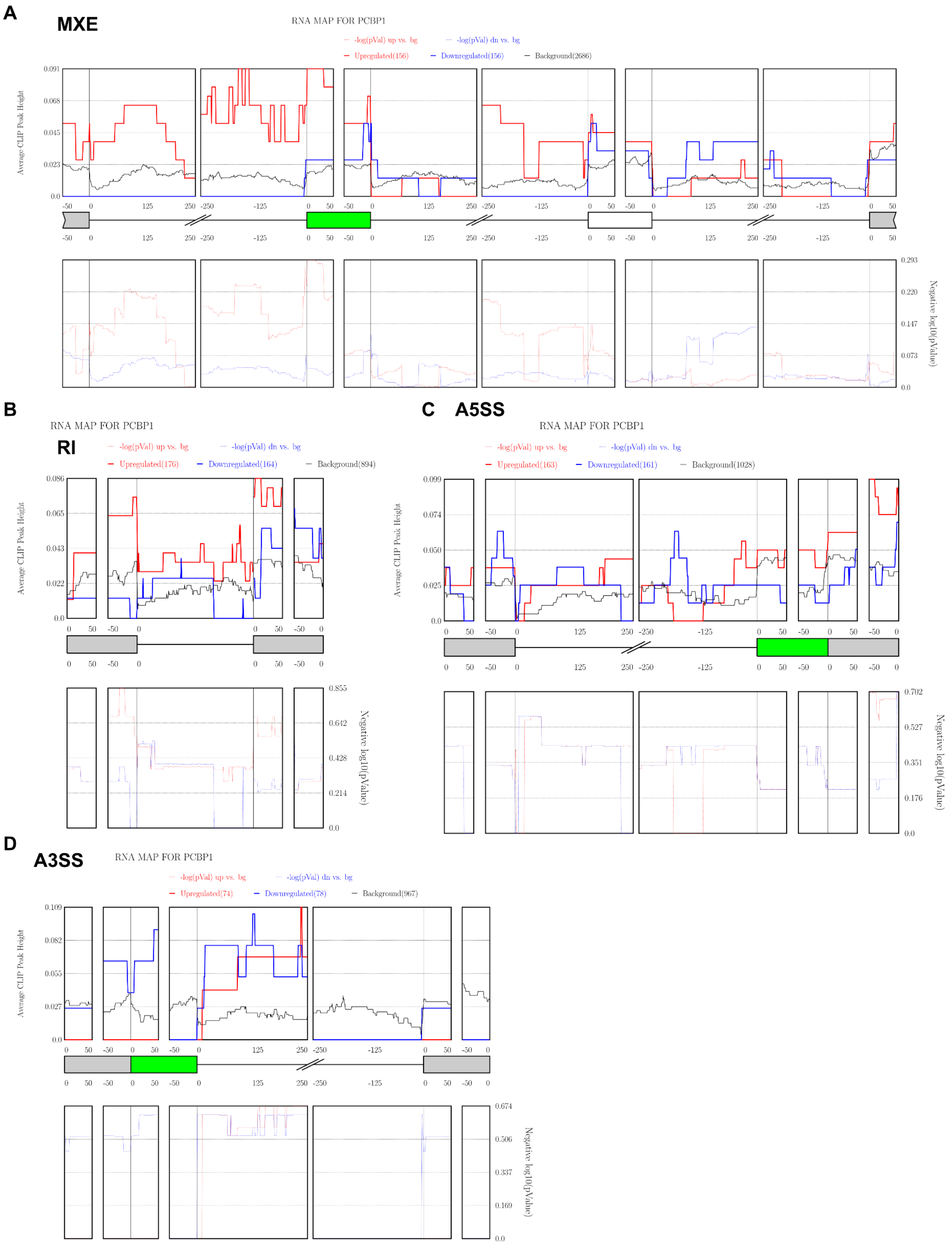
A
MXE
RNA MAP FOR PCBP1
-log(pVal) up vs. bg
-log(pVal) dn vs. bg
Upregulated(156)
Downregulated(156)
Background(2686)
Average CLIP Peak Height
Negative log10(pValue)
B
RNA MAP FOR PCBP1
RI
-log(pVal) up vs. bg
-log(pVal) dn vs. bg
Upregulated(176)
Downregulated(164)
Background(894)
Average CLIP Peak Height
Negative log10(pValue)
C
A5SS
RNA MAP FOR PCBP1
-log(pVal) up vs. bg
-log(pVal) dn vs. bg
Upregulated(163)
Downregulated(161)
Background(1028)
Average CLIP Peak Height
Negative log10(pValue)
D
A3SS
RNA MAP FOR PCBP1
-log(pVal) up vs. bg
-log(pVal) dn vs. bg
Upregulated(74)
Downregulated(78)
Background(967)
Average CLIP Peak Height
Negative log10(pValue)

**Supplemental Figure 5. rMAPS2 analysis revealed minimal enrichment for PCBP1 eCLIP peaks at MXE, RI, A5SS and A3SS events detected in the Pcbp1-cKO heart.** (A-D) RNA maps for PCBP1 were analyzed by rMAPS2, and the PCBP1 eCLIP peak enrichment were visualized on each alternative spliced pattern that was detected to be differentially regulated by Pcbp1-cKO.

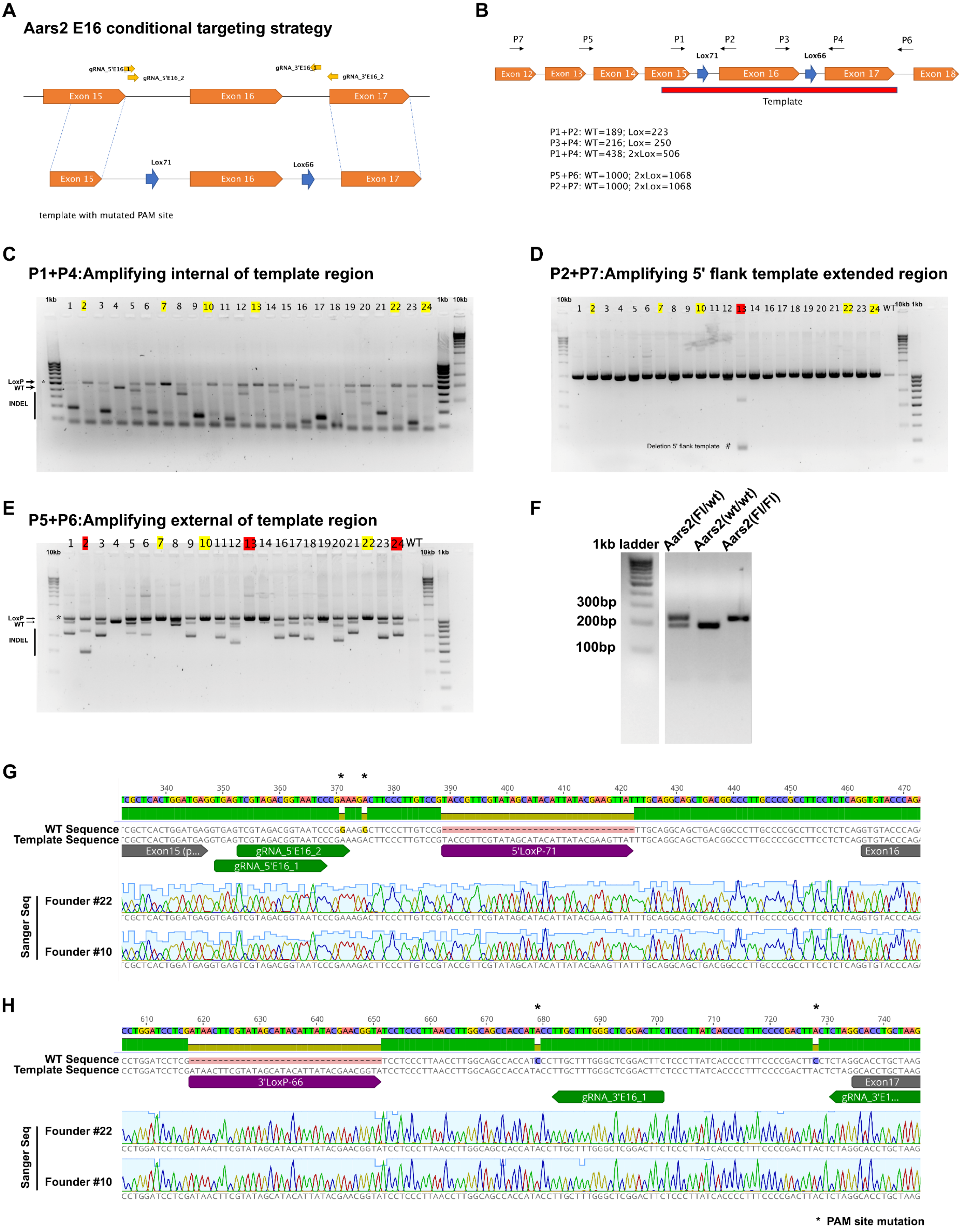
A
Aars2 E16 conditional targeting strategy
gRNA_5'E16_1
gRNA_5'E16_2
gRNA_3'E16_1
gRNA_3'E16_2
Exon 15
Exon 16
Exon 17
Lox71
Lox66
template with mutated PAM site
B
P7
P5
P1
P2
P3
P4
P6
Lox71
Lox66
Exon 12
Exon 13
Exon 14
Exon 15
Exon 16
Exon 17
Exon 18
Template
P1+P2: WT=189; Lox=223
P3+P4: WT=216; Lox= 250
P1+P4: WT=438; 2xLox=506
P5+P6: WT=1000; 2xLox=1068
P2+P7: WT=1000; 2xLox=1068
C
P1+P4:Amplifying internal of template region
1kb 1 2 3 4 5 6 7 8 9 10 11 12 13 14 15 16 17 18 19 20 21 22 23 24 1kb 10kb
LoxP
WT
INDEL
*
D
P2+P7:Amplifying 5' flank template extended region
10kb 1 2 3 4 5 6 7 8 9 10 11 12 13 14 16 17 18 19 20 21 22 23 24 WT 10kb 1kb
Deletion 5' flank template #
E
P5+P6:Amplifying external of template region
1 2 3 4 5 6 7 8 9 10 11 12 13 14 16 17 18 19 20 21 22 23 24 WT
10kb
10kb 1kb
LoxP
WT
INDEL
*
F
1kb ladder
Aars2(Fl/wt)
Aars2(wt/wt)
Aars2(Fl/Fl)
300bp
200bp
100bp
G
WT Sequence
Template Sequence
Exon15 (p...
gRNA_5'E16_2
gRNA_5'E16_1
5'LoxP-71
Exon16
Sanger Seq
Founder #22
Founder #10
H
WT Sequence
Template Sequence
3'LoxP-66
Exon17
gRNA_3'E16_1
gRNA_3'E1...
Sanger Seq
Founder #22
Founder #10
* PAM site mutation

**Supplemental Figure 6. The generation of Aars2 loxP flanked allele for exon-16 targeting.** (A) Two loxP sites flanking exon-16 of Aars2 were introduced by homologous recombination using CRISPR/Cas9 strategy to incorporate a homologous DNA template with mutated PAM sites. Two combination pairs of gRNAs from 5' and 3' of the template were utilized. (B) Genotyping strategy to amplify different genomic fragment internal or external of the insertion were designed to confirm correct insertion of the template. 24 viable pups from two rounds of injections were recovered, and subsequently screened. (C-E) Using PCR to amplify internal or external of the template insertion, we detected founders with various sizes of indels, as well as founders with correct insertions (highlight in yellow). (F) Two Aars2-loxP harboring lines were selected and backcrossed for 4 generations. Representative of the loxP genotyping were shown using F1+F2 primer combinations. (G-H) PCR products flanking both loxP sites from the genomic DNA of these founders were subsequently TA-cloned, and at least 5 colonies were Sanger sequenced to confirm correct loxP integration in each allele. Representative of 5' loxP site region (G) and the 3' loxP site region (H) are shown. With the correct loxP insertion, we also observed the correctly incorporated mutated PAM site engineered within the DNA template.

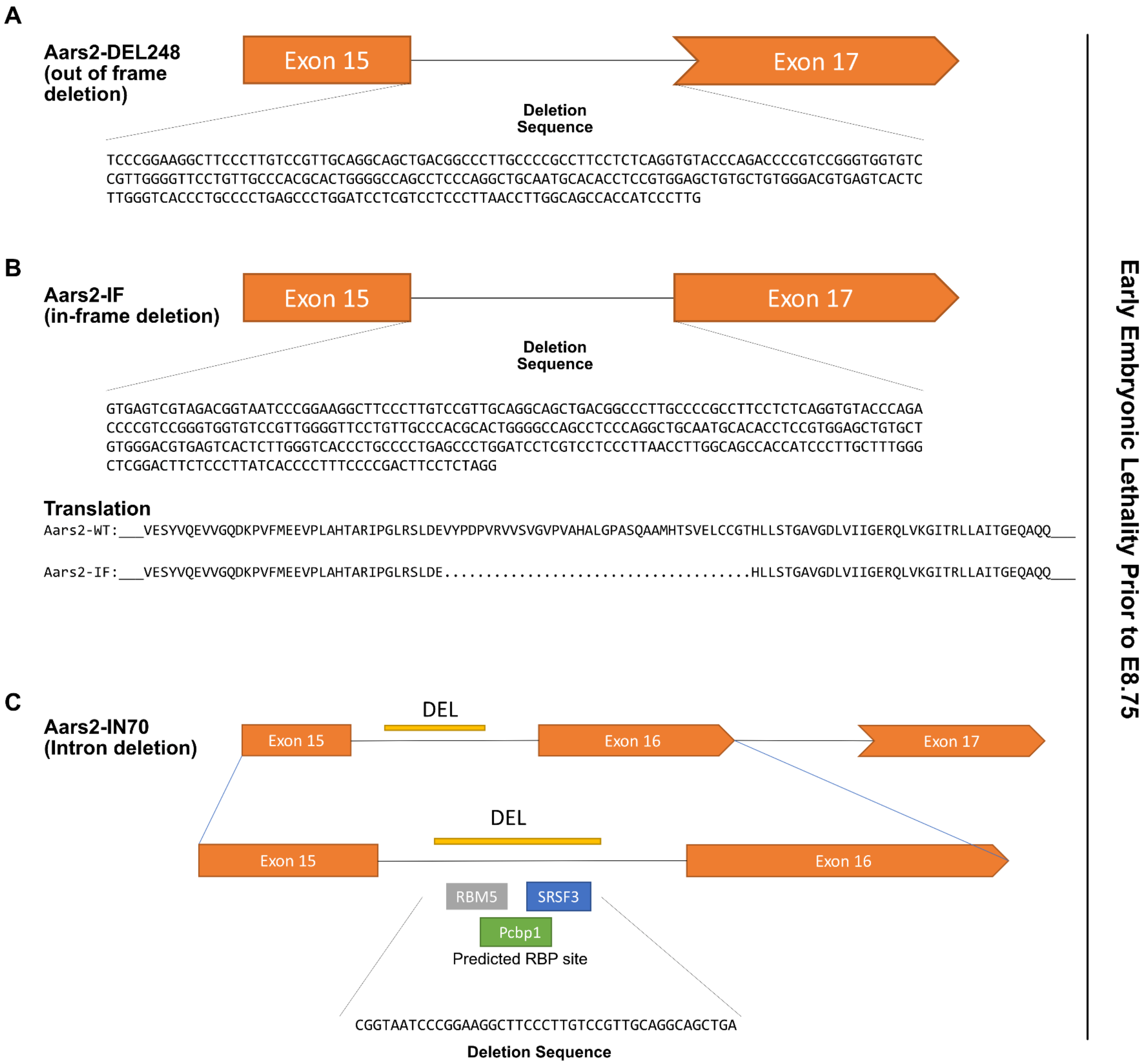
A
Aars2-DEL248
(out of frame
deletion)
Exon 15
Exon 17
Deletion
Sequence
TCCCGGAAGGCTTCCCTTGTCCGTTGCAGGCAGCTGACGGCCCTTGCCCCGCCTTCCTCTCAGGTGTACCCAGACCCCGTCCGGGTGGTGTC
CGTTGGGGTTCCTGTTGCCCACGCACTGGGGCCAGCCTCCCAGGCTGCAATGCACACCTCCGTGGAGCTGTGCTGTGGGACGTGAGTCACTC
TTGGGTCACCCTGCCCCTGAGCCCTGGATCCTCGTCCTCCCTTAACCTTGGCAGCCACCATCCCTTG
B
Aars2-IF
(in-frame deletion)
Exon 15
Exon 17
Deletion
Sequence
GTGAGTCGTAGACGGTAATCCCGGAAGGCTTCCCTTGTCCGTTGCAGGCAGCTGACGGCCCTTGCCCCGCCTTCCTCTCAGGTGTACCCAGA
CCCCGTCCGGGTGGTGTCCGTTGGGGTTCCTGTTGCCCACGCACTGGGGCCAGCCTCCCAGGCTGCAATGCACACCTCCGTGGAGCTGTGCT
GTGGGACGTGAGTCACTCTTGGGTCACCCTGCCCCTGAGCCCTGGATCCTCGTCCTCCCTTAACCTTGGCAGCCACCATCCCTTGCTTTGGG
CTCGGACTTCTCCCTTATCACCCCTTTCCCCGACTTCCTCTAGG
Translation
Aars2-WT:___VESYVQEVVGQDKPVFMEEVPLAHTARIPGLRSLDEVYPDPVRVVSVGVPVAHALGPASQAAMHTSVELCCGTHLLSTGAVGDLVIIGERQLVKGITRLLAITGEQAQQ___
Aars2-IF:___VESYVQEVVGQDKPVFMEEVPLAHTARIPGLRSLDE.....................................HLLSTGAVGDLVIIGERQLVKGITRLLAITGEQAQQ___
C
Aars2-IN70
(Intron deletion)
DEL
Exon 15
Exon 16
Exon 17
DEL
Exon 15
Exon 16
RBM5
SRSF3
Pcbp1
Predicted RBP site
CGGTAATCCCGGAAGGCTTCCCTTGTCCGTTGCAGGCAGCTGA
Deletion Sequence
Early Embryonic Lethality Prior to E8.75

**Supplemental Figure 7. Homozygous germline deletion of Aars2 exon-16 leads to early embryonic lethality.** Through screening the founders, we selected three germline deletion mutants to carry through back-crossing before crossing to homozygosity. (A) Aars2-DEL248 mutant carries a deletion region removing exon 16, and is predicted to place the downstream transcript out of frame. (B) Aars2-IF mutant carries a deletion that removes the intron flanking exon 16 as well as exon 16, and one extra nucleotide on exon 17 that is predicted to produce an in-frame deletion of exon 16. (C) Aars2-IN70 mutant carries a 43 bp deletion at the intron between exon 15 and exon 16, and the deleted region contains predicted binding sites for RBM5, SRSF3, and PCBP1. Heterozygous animals of all three deletion mutants were inter-crossed independently to generate homozygous mutants, but no homozygous mutants were recovered at E8.75, suggesting early embryonic lethality.

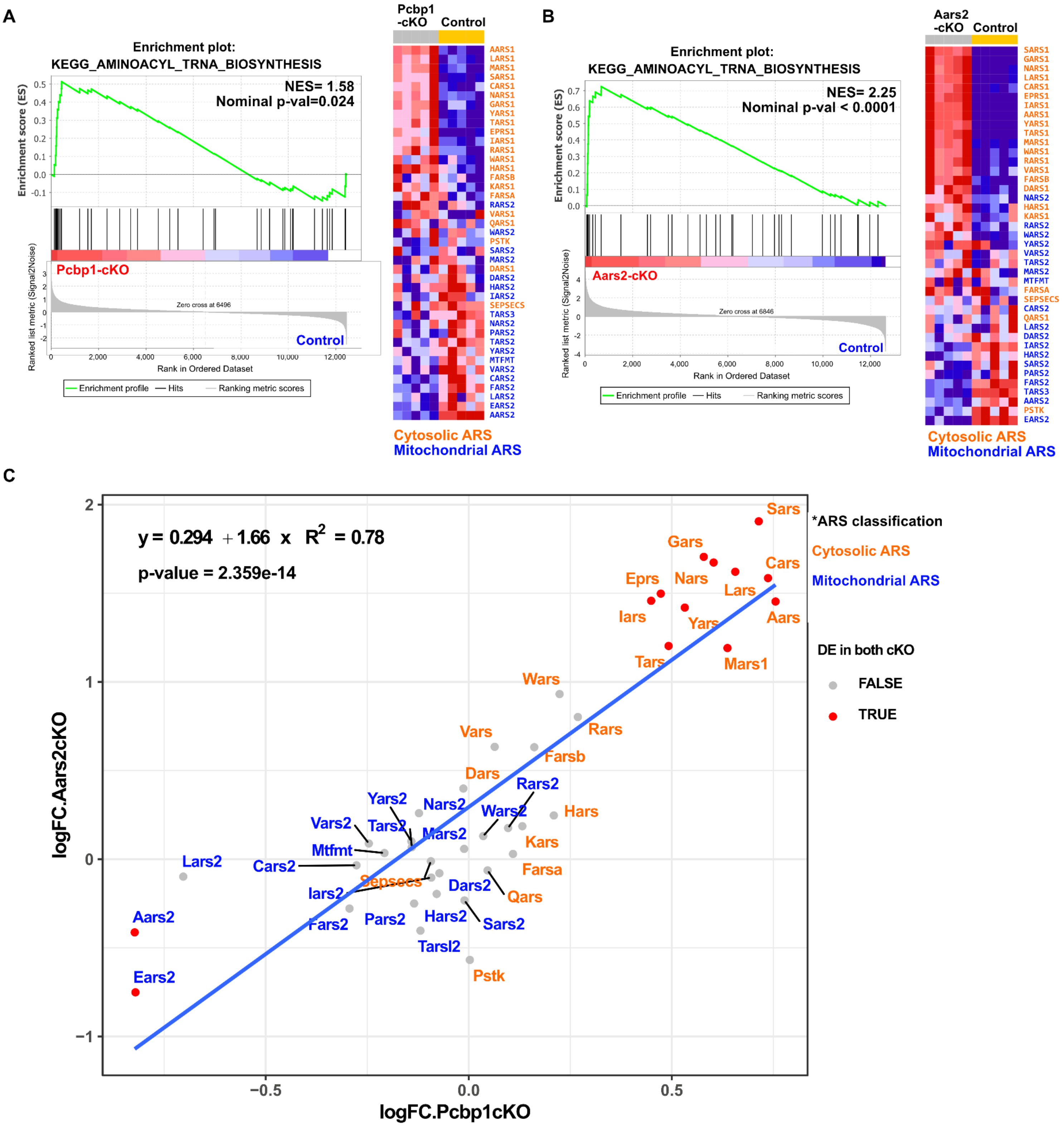
A
Pcbp1 -cKO
Control
Enrichment plot:
KEGG_AMINOACYL_TRNA_BIOSYNTHESIS
NES= 1.58
Nominal p-val=0.024
Enrichment score (ES)
Pcbp1-cKO
Control
Zero cross at 6496
Ranked list metric (Signal2Noise)
Rank in Ordered Dataset
Enrichment profile
Hits
Ranking metric scores
AARS1 LARS1 MARS1 SARS1 CARS1 NARS1 GARS1 YARS1 TARS1 EPRS1 IARS1 RARS1 WARS1 HARS1 FARSB KARS1 FARSA RARS2 VARS1 QARS1 WARS2 PSTK SARS2 MARS2 DARS1 DARS2 HARS2 IARS2 SEPSECS TARS3 NARS2 PARS2 TARS2 YARS2 MTFMT VARS2 CARS2 FARS2 LARS2 EARS2 AARS2
Cytosolic ARS
Mitochondrial ARS
B
Aars2 -cKO
Control
Enrichment plot:
KEGG_AMINOACYL_TRNA_BIOSYNTHESIS
NES= 2.25
Nominal p-val < 0.0001
Enrichment score (ES)
Aars2-cKO
Control
Zero cross at 6846
Ranked list metric (Signal2Noise)
Rank in Ordered Dataset
Enrichment profile
Hits
Ranking metric scores
SARS1 GARS1 NARS1 LARS1 CARS1 EPRS1 IARS1 AARS1 YARS1 TARS1 MARS1 WARS1 RARS1 VARS1 FARSB DARS1 NARS2 HARS1 KARS1 RARS2 WARS2 YARS2 VARS2 TARS2 MARS2 MTFMT FARSA SEPSECS CARS2 QARS1 LARS2 DARS2 IARS2 HARS2 SARS2 PARS2 FARS2 TARS3 AARS2 PSTK EARS2
Cytosolic ARS
Mitochondrial ARS
C
y = 0.294 + 1.66 x R2 = 0.78
p-value = 2.359e-14
*ARS classification
Cytosolic ARS
Mitochondrial ARS
DE in both cKO
FALSE
TRUE
Sars Gars Cars Eprs Nars Lars Iars Yars Aars Tars Mars1 Wars Rars Vars Farsb Dars Rars2 Hars Yars2 Nars2 Wars2 Vars2 Tars2 Mars2 Kars Mtfmt Lars2 Cars2 Sepsecs Farsa Iars2 Dars2 Qars Aars2 Fars2 Pars2 Hars2 Sars2 Tarsl2 Ears2 Pstk
logFC.Aars2cKO
logFC.Pcbp1cKO

**Supplemental Figure 8. Concordant up-regulation of cytosolic ARS in both Pcbp1-cKO and Aars2-cKO transcriptome.** KEGG gene-sets of aminoacyl tRNA biosynthesis are positively enriched in both Pcbp1-cKO (A) and Aars2-cKO (B). The cytosolic Aminoacyl-tRNA Synthetase (ARS) (label in orange) are collectively up-regulated in both mutants. (C) Plotting the logFC of cytosolic ARS and mitochondrial ARS from both Pcbp1-cKO and Aars2-cKO with linear regression revealed a concordant regulation for the cytosolic ARS, with a large subset of 11 cytosolic ARS up-regulated in both mutants, while only 2 mitochondrial ARS genes (Aars2 and Ears2) were downregulated in both mutants.
