## Supplemental Tables for "PCBP1 regulates alternative splicing of AARS2 in congenital cardiomyopathy"

**Supplementary Table 1. Echocardiography examination of Pcbp1-cKO mice and their littermate controls immediately after birth (P0).**

| | Ctnt-Cre(-) Cntl (N=21) | Pcbp1-cHet (N=7) | Pcbp1-cKO (N=11) |
|---|---|---|---|
| **EF (%)** | 81.5 ± 7.99 | 83.0 ± 6.95 | 67.1 ± 15.4 ** |
| **FS (%)** | 47.5 ± 8.85 | 48.7 ± 6.94 | 35.3 ± 11.1 ** |
| **LVID;d (mm)** | 1.31 ± 0.129 | 1.34 ± 0.190 | 1.34 ± 0.294 |
| **LVID;s (mm)** | 0.691 ± 0.142 | 0.698 ± 0.186 | 0.880 ± 0.284 * |
| **LVPW.d (mm)** | 0.396 ± 0.0551 | 0.399 ± 0.0710 | 0.412 ± 0.0900 |
| **LVPW.s (mm)** | 0.622 ± 0.0979 | 0.622 ± 0.0660 | 0.555 ± 0.0847 |
| **LV.Mass (mg)** | 7.92 ± 1.46 | 7.65 ± 1.53 | 8.07 ± 3.01 |
| **LV.Mass(corrected) (mg)** | 6.34 ± 1.17 | 6.12 ± 1.22 | 6.45 ± 2.41 |
| **LV.Vol;d (ul)** | 4.35 ± 1.06 | 4.67 ± 1.69 | 4.87 ± 2.44 |
| **LV.Vol;s (ul)** | 0.819 ± 0.445 | 0.881 ± 0.634 | 1.76 ± 1.26 ** |
| **Heart.Rate (BPM)** | 314 ± 72.4 | 410 ± 47.1 ** | 277 ± 46.0 |

One-way ANOVA follow by Turkey Post-hoc test was used for comparison of Pcbp1-cKO or Pcbp1-cHet with Ctnt0Cre(-) Control. Corrected p-value is reported as follow: * $p < 0.05$, ** $p < 0.01$.

**Supplementary Table 2. Echocardiography examination of Pcbp1-cKO mice and their littermate controls at P1.**

| | Ctnt-Cre(-) Cntl (N=12) | Pcbp1-cHet (N=6) | Pcbp1-cKO (N=6) |
|---|---|---|---|
| **EF (%)** | 87.3 ± 4.43 | 88.1 ± 2.54 | 68.2 ± 14.5 *** |
| **FS (%)** | 54.0 ± 5.98 | 54.3 ± 3.19 | 36.4 ± 11.6 *** |
| **LVID;d (mm)** | 1.43 ± 0.171 | 1.26 ± 0.236 | 1.55 ± 0.298 |
| **LVID;s (mm)** | 0.662 ± 0.149 | 0.579 ± 0.143 | 1.01 ± 0.348 ** |
| **LVPW.d (mm)** | 0.469 ± 0.0795 | 0.478 ± 0.116 | 0.408 ± 0.0853 |
| **LVPW.s (mm)** | 0.732 ± 0.0858 | 0.711 ± 0.114 | 0.572 ± 0.068 ** |
| **LV.Mass (mg)** | 10.7 ± 2.66 | 8.80 ± 1.44 | 9.84 ± 4.39 |
| **LV.Mass(corrected) (mg)** | 8.53 ± 2.13 | 7.04 ± 1.15 | 7.87 ± 3.51 |
| **LV.Vol;d (ul)** | 5.44 ± 1.63 | 4.03 ± 2.15 | 7.05 ± 3.51 |
| **LV.Vol;s (ul)** | 0.737 ± 0.427 | 0.514 ± 0.380 | 2.57 ± 2.31 * |
| **Heart.Rate (BPM)** | 408 ± 54.0 | 418 ± 56.4 | 323 ± 68.6 * |

One-way ANOVA follow by Turkey Post-hoc test was used for comparison of Pcbp1-cKO or Pcbp1-cHet with Ctnt0Cre(-) Control. Corrected p-value is reported as follow: * p < 0.05, ** p< 0.01, *** p < 0.001.

**Supplementary Table 3. Echocardiography examination of Pcbp1-cKO mice and their littermate controls at P2.**

| | Ctnt-Cre(-) Cntl (N=10) | Pcbp1-cHet (N=5) | Pcbp1-cKO (N=4) |
|---|---|---|---|
| **EF (%)** | 85.5 ± 5.03 | 83.9 ± 5.26 | 61.8 ± 14.4 *** |
| **FS (%)** | 52.1 ± 5.97 | 50.0 ± 5.78 | 31.6 ± 9.55 *** |
| **LVID;d (mm)** | 1.73 ± 0.216 | 1.66 ± 0.157 | 1.85 ± 0.218 |
| **LVID;s (mm)** | 0.838 ± 0.198 | 0.833 ± 0.166 | 1.28 ± 0.329 * |
| **LVPW.d (mm)** | 0.502 ± 0.0429 | 0.450 ± 0.0292 | 0.410 ± 0.112 |
| **LVPW.s (mm)** | 0.771 ± 0.0530 | 0.695 ± 0.0572 | 0.602 ± 0.121 ** |
| **LV.Mass (mg)** | 16.2 ± 3.42 | 14.0 ± 3.45 | 14.1 ± 6.40 |
| **LV.Mass(corrected) (mg)** | 13.0 ± 2.74 | 11.2 ± 2.76 | 11.2 ± 5.12 |
| **LV.Vol;d (ul)** | 9.01 ± 2.87 | 7.95 ± 2.04 | 10.6 ± 3.22 |
| **LV.Vol;s (ul)** | 1.42 ± 0.884 | 1.35 ± 0.796 | 4.41 ± 2.93 ** |
| **Heart.Rate (BPM)** | 424 ± 44.8 | 447 ± 61.2 | 311 ± 86.0 * |

One-way ANOVA follow by Turkey Post-hoc test was used for comparison of Pcbp1-cKO or Pcbp1-cHet with Ctnt0Cre(-) Control. Corrected p-value is reported as follow: * $p < 0.05$, ** $p< 0.01$, *** $p < 0.001$.

**Supplementary Table 4. Aars2 Exon 16 frameshift and translation summary.**

**1. Aars2 WT cDNA (Exon-16 highlighted)**

atggcggtggcgttggctgccgcagccggtaagctgcggcgagccattgggaggtcgtgcccatggc
agcctttctcaaccgagcccggtccaccccacggagcggccgtgcgggacgccttcctgagcttctt
ccgagatcgccacggccaccggctcgtgccctccgctaccgtgaggccgcgcggcgaccccagcctg
cttttcgtcaatgcaggcatgaaccagttcaagcccatcttcctgggcacagtggatccacgaagtg
agatggcaggcttccgacgtgtagttaatagccagaagtgtgtcagggctggaggacgccataacga
cctggaggatgtggggcgtgatctctctcatcatacgtttttcgagatgcttggcaactgggctttc
gggggtgaatattttaaggaggaagcttgtagcatggcctgggaacttctgactcaagtctacggga
tccctgaggacaggttgtgggtctcctacttcagtggtgactcccagacaggactggacccagacct
ggagaccagagacatctggctaagcttgggagtgcctgccagccgtgtgctctccttcggaccacaa
gagaacttctgggagatgggagacactgggccttgtgggccttgtactgagatccactatgacctgg
ctgggggcgtgggaagcccccagctggtagagctttggaatctggtcttcatgcaacactacagaga
ggcagatggaagcctgcagctgctgccgcagcgacatgtagacacaggaatgggcctggaaaggctg
gtggccgtcctgcaaggcaaacgttccacctacgacaccgacctcttctctccactgctcgacgcca
tacaccagagctgcggggctccccttactctggccgggtaggggcagcagatgagggacggataga
cacagcgtaccgagtggtagccgatcacatccgcacactcagtgtctgcattgctgacggtgtctcc
ccggggatgtcaggtgccccgctagttctccgtcggattctccgccgagctgtgcgctattccacag
aggtcttgcaggcacctcctggctttctaggcagcctggtgccagtggtggtagagacactgggatc
tgcttatccagaacttgagaagaactcagtcaagatagccagcctggtgtcagaggatgaggcagcc
ttcctggcctccctgcagcggggccggcggatcatcgatcgcaccgtcaagcgcctgggcccttctg
atttgttccctgccgaagtggcctggtcattgtcgctgtctgggaatctggggattcccctggacct
ggtggagctgatgctggaggagaagggggtgaagctggacacggcaggactggagcagctagcccag
aaggaggctcagcaccgggcccagcaagcagaggcagatcaggaggaccgattgtgtcttgatgtcc
acgcactggaggagctgcaccgtcaaggcatacccacaactgatgacagccccaagtataactacac
tcttcaccccaacggggattatgagtttggcctctgtgaggcccgggtgttacagctgtattcagag
actgggacagccgtggcctccgtgggagcaggccagcgctgtggcctcctattagatagaaccaact
tctacgctgaacaagggggtcaggcttcagaccgaggctaccttgtccgtacagggcagcaggacat
gttgttccctgtggctggggctcagctctgtgggggttttatcctgcatgaggccatggctcccgag
cgcctacaggtgggggatcaagtgcagctgtatgtagataaggcctggcgaatgggatgcatggtga
agcacacggccacccacctgctgagctgggcccttcggcagaccctcggaccgaccaccgagcagcg
gggctcccatctcaaccccgagcggctgcgctttgacgtggccacccagaccctactaaccacagag
cagctacggacagtagagagctacgtgcaggaagtcgtggggcaggataagcctgtgttcatggagg
aagtgcccctggcacacactgcccgcatccccggccttcgctcactggatgaggtgtacccagaccc
cgtccgggtggtgtccgttggggttcctgttgcccacgcactggggccagcctcccaggctgcaatg
cacacctccgtggagctgtgctgtgggacgcacctgctaagtaccggggccgtgggagacttggtga
tcattggggaacggcagctggtcaagggcatcactcgcctactggccatcactggggagcaggccca
acaggcccgagaggtaggccagagcctgtctcaggaggtggaagcagccagcgagcgactaagtcag
ggcagccgagacctgccggaagctcaccggctatctaaggacataggacgtctcaccgaggtagcag
agtctgctgtgatacctcagtggcaacggcaggagctgcagaccacactgaagatgctacagcgccg
tgccaacactgccatccggaaactggagaagggccaggctacagagaaatcccaggagctgttgaaa
cgacactcggaggggcctctgattgtggacactgtctctgccgagtccctctcagtgctggtaaagg
tcgtgcggcagctgtgcaaacaggcccctagcatatccgtgctgctgctcagcccccagcccactgg
cagtgtcctgtgtgcctgccaggtggcccaggatgccacgcccaccttcactgctgaagcctgggca
ctggctgtgtgcagccacatgggaggcaaggcgtggggctccagagtggtagctcaggggactggac
acacggctgacctggaggccgccctcgggacagcccgagcttacgcactcagccagctc

**2. Aars2 Exon-16 skipping cDNA**

atggcggtggcgttggctgccgcagccggtaagctgcggcgagccattgggaggtcgtgcccatggc
agcctttctcaaccgagcccggtccaccccacggagcggccgtgcgggacgccttcctgagcttctt
ccgagatcgccacggccaccggctcgtgccctccgctaccgtgaggccgcgcggcgaccccagcctg
cttttcgtcaatgcaggcatgaaccagttcaagcccatcttcctgggcacagtggatccacgaagtg
agatggcaggcttccgacgtgtagttaatagccagaagtgtgtcagggctggaggacgccataacga
cctggaggatgtggggcgtgatctctctcatcatacgtttttcgagatgcttggcaactgggctttc
gggggtgaatattttaaggaggaagcttgtagcatggcctgggaacttctgactcaagtctacggga
tccctgaggacaggttgtgggtctcctacttcagtggtgactcccagacaggactggacccagacct
ggagaccagagacatctggctaagcttgggagtgcctgccagccgtgtgctctccttcggaccacaa
gagaacttctgggagatgggagacactgggccttgtgggccttgtactgagatccactatgacctgg
ctgggggcgtgggaagcccccagctggtagagctttggaatctggtcttcatgcaacactacagaga
ggcagatggaagcctgcagctgctgccgcagcgacatgtagacacaggaatgggcctggaaaggctg
gtggccgtcctgcaaggcaaacgttccacctacgacaccgacctcttctctccactgctcgacgcca
tacaccagagctgcggggctccccttactctggccgggtaggggcagcagatgagggacggataga
cacagcgtaccgagtggtagccgatcacatccgcacactcagtgtctgcattgctgacggtgtctcc
ccggggatgtcaggtgccccgctagttctccgtcggattctccgccgagctgtgcgctattccacag
aggtcttgcaggcacctcctggctttctaggcagcctggtgccagtggtggtagagacactgggatc
tgcttatccagaacttgagaagaactcagtcaagatagccagcctggtgtcagaggatgaggcagcc
ttcctggcctccctgcagcggggccggcggatcatcgatcgcaccgtcaagcgcctgggcccttctg
atttgttccctgccgaagtggcctggtcattgtcgctgtctgggaatctggggattcccctggacct
ggtggagctgatgctggaggagaagggggtgaagctggacacggcaggactggagcagctagcccag
aaggaggctcagcaccgggcccagcaagcagaggcagatcaggaggaccgattgtgtcttgatgtcc
acgcactggaggagctgcaccgtcaaggcatacccacaactgatgacagccccaagtataactacac
tcttcaccccaacggggattatgagtttggcctctgtgaggcccgggtgttacagctgtattcagag
actgggacagccgtggcctccgtgggagcaggccagcgctgtggcctcctattagatagaaccaact
tctacgctgaacaagggggtcaggcttcagaccgaggctaccttgtccgtacagggcagcaggacat
gttgttccctgtggctggggctcagctctgtgggggttttatcctgcatgaggccatggctcccgag
cgcctacaggtgggggatcaagtgcagctgtatgtagataaggcctggcgaatgggatgcatggtga
agcacacggccacccacctgctgagctgggcccttcggcagaccctcggaccgaccaccgagcagcg
gggctcccatctcaaccccgagcggctgcgctttgacgtggccacccagaccctactaaccacagag
cagctacggacagtagagagctacgtgcaggaagtcgtggggcaggataagcctgtgttcatggagg
aagtgcccctggcacacactgcccgcatccccggccttcgctcactggatgaggcacctgctaagta
ccggggccgtgggagacttggtgatcattggggaacggcagctggtcaagggcatcactcgcctact
ggccatcactggggagcaggcccaacaggcccgagaggtaggccagagcctgtctcaggaggtggaa
gcagccagcgagcgactaagtcagggcagccgagacctgccggaagctcaccggctatctaaggaca
taggacgtctcaccgaggtagcagagtctgctgtgatacctcagtggcaacggcaggagctgcagac
cacactgaagatgctacagcgccgtgccaacactgccatccggaaactggagaagggccaggctaca
gagaaatcccaggagctgttgaaacgacactcggaggggcctctgattgtggacactgtctctgccg
agtccctctcagtgctggtaaaggtcgtgcggcagctgtgcaaacaggcccctagcatatccgtgct
gctgctcagcccccagcccactggcagtgtcctgtgtgcctgccaggtggcccaggatgccacgccc
accttcactgctgaagcctgggcactggctgtgtgcagccacatgggaggcaaggcgtggggctcca
gagtggtagctcaggggactggacacacggctgacctggaggccgccctcgggacagcccgagctta
cgcactcagccagctc

**3. Aars2 WT cDNA Translation**

MAVALAAAAGKLRRAIGRSCPWQPFSTEPGPPHGAAVRDAFLSFFRDRHGHRLVPSATVRPRGDPSL
LFVNAGMNQFKPIFLGTVDPRSEMAGFRRVVNSQKCVRAGGRHNDLEDVGRDLSHHTFFEMLGNWAF
GGEYFKEEACSMAWELLTQVYGIPEDRLWVSYFSGDSQTGLDPDLETRDIWLSLGVPASRVLSFGPQ
ENFWEMGDTGPCGPCTEIHYDLAGGVGSPQLVELWNLVFMQHYREADGSLQLLPQRHVDTGMGLERL
VAVLQGKRSTYDTDLFSPLLDAIHQSCGAPPYSGRVGAADEGRIDTAYRVVADHIRTLSVCIADGVS
PGMSGAPLVLRRILRRAVRYSTEVLQAPPGFLGSLVPVVVETLGSAYPELEKNSVKIASLVSEDEAA
FLASLQRGRRIIDRTVKRLGPSDLFPAEVAWSLSLSGNLGIPLDLVELMLEEKGVKLDTAGLEQLAQ
KEAQHRAQQAEADQEDRLCLDVHALEELHRQGIPTTDDSPKYNYTLHPNGDYEFGLCEARVLQLYSE
TGTAVASVGAGQRCGLLLDRTNFYAEQGGQASDRGYLVRTGQQDMLFPVAGAQLCGGFILHEAMAPE
RLQVGDQVQLYVDKAWRMGCMVKHTATHLLSWALRQTLGPTTEQRGSHLNPERLRFDVATQTLLTTE
QLRTVESYVQEVVGQDKPVFMEEVPLAHTARIPGLRSLDEVYPDPVRVVSVGVPVAHALGPASQAAM
HTSVELCCGTHLLSTGAVGDLVIIGERQLVKGITRLLAITGEQAQQAREVGQSLSQEVEAASERLSQ
GSRDLPEAHRLSKDIGRLTEVAESAVIPQWQRQELQTTLKMLQRRANTAIRKLEKGQATEKSQELLK
RHSEGPLIVDTVSAESLSVLVKVVRQLCKQAPSISVLLLSPQPTGSVLCACQVAQDATPTFTAEAWA
LAVCSHMGGKAWGSRVVAQGTGHTADLEAALGTARAYALSQL

**4. Aars2 Exon-16 skipping cDNA Translation (* premature stop, frame-shift)**

MAVALAAAAGKLRRAIGRSCPWQPFSTEPGPPHGAAVRDAFLSFFRDRHGHRLVPSATVRPRGDPSL
LFVNAGMNQFKPIFLGTVDPRSEMAGFRRVVNSQKCVRAGGRHNDLEDVGRDLSHHTFFEMLGNWAF
GGEYFKEEACSMAWELLTQVYGIPEDRLWVSYFSGDSQTGLDPDLETRDIWLSLGVPASRVLSFGPQ
ENFWEMGDTGPCGPCTEIHYDLAGGVGSPQLVELWNLVFMQHYREADGSLQLLPQRHVDTGMGLERL
VAVLQGKRSTYDTDLFSPLLDAIHQSCGAPPYSGRVGAADEGRIDTAYRVVADHIRTLSVCIADGVS
PGMSGAPLVLRRILRRAVRYSTEVLQAPPGFLGSLVPVVVETLGSAYPELEKNSVKIASLVSEDEAA
FLASLQRGRRIIDRTVKRLGPSDLFPAEVAWSLSLSGNLGIPLDLVELMLEEKGVKLDTAGLEQLAQ
KEAQHRAQQAEADQEDRLCLDVHALEELHRQGIPTTDDSPKYNYTLHPNGDYEFGLCEARVLQLYSE
TGTAVASVGAGQRCGLLLDRTNFYAEQGGQASDRGYLVRTGQQDMLFPVAGAQLCGGFILHEAMAPE
RLQVGDQVQLYVDKAWRMGCMVKHTATHLLSWALRQTLGPTTEQRGSHLNPERLRFDVATQTLLTTE
QLRTVESYVQEVVGQDKPVFMEEVPLAHTARIPGLRSLDE**APAKYRGRGRLGDHWGTAAGQGHHSPT**
**GHHWGAGPTGPRGRPEPVSGGGSSQRATKSGQPRPAGSSPAI***GHRTSHRGSRVCCDTSVATAGAAD
HTEDATAPCQHCHPETGEGPGYREIPGAVETTLGGASDCGHCLCRVPLSAGKGRAAAVQTGP*HIRA
AAQPPAHWQCPVCLPGGPGCHAHLHC*SLGTGCVQPHGRQGVGLQSGSSGDWTHG-
PGGRPRDSPSLRTQPA

### Supplemental Table 5. qRT-PCR primer sequences.

| Gene | Forward | Reverse | Reactivity (M=Mouse, R=Rat, H=Human) |
|---|---|---|---|
| Nppa | TACAGTGCGGTGTCCAACACAG | TGCTTCCTCAGTCTGCTCACTC | M |
| 18S | CATTCGAACGTCTGCCCTAT | GTTTCTCAGGCTCCCTCTCC | MRH |
| Pcbp1 | GGACAACACACCATTTCTCCGC | AGCCTTTCACCTCTGGAGAGCT | MRH |
| Pcbp2 | CTTTGGCTGGACCGACTAATGC | TAGCAGGAACCACCAGCCGTAA | MR |
| Notch2 | AGACTGGCGACTTCACTTTC | CAGCGGCAATTGTAGGTATTG | M |
| Jag1 | TCGGGAGGCAAATTCACC | GCCGTCACTACAGATACACTTG | M |
| Hey2 | CAAGGATCTGCCAAGTTAGAAAAG | TGTCAAGCACTCTCGGAATC | M |
| Atf4 | GGTTCTCCAGCGACAAGG | GCATCGAAGTCAAACTCTTTCAG | M |
| Ddit3 | GCACCTATATCTCATCCCCAG | TGCGTGTGACCTCTGTTG | M |
| Asns | AATGCAGCCGATAAGAGTGAG | CCTTTGTCATAGAGGTGGAGG | M |
| Hprt | TGGCCCTCTGTGTGCTCAA | TGATCATTACAGTAGCTCTTCAGTCTGA | M |
| Aars2 (exon 14-15) | TCCCATCTCAACCCCGAG | TCCATGAACACAGGCTTATCC | M |
| Aars2 (exon 15 skip to 17) | CACTGGATGAGGCACCTG | GCCAGTAGGCGAGTGATG | MR |
| Aars2 (exon 16-17) | GGTTCCTGTTGCCCACG | GAGTGATGCCCTTGACCAG | M |

**Supplemental Table 6. Genotyping primer sequences.**

| | |
|---|---|
| (1+2:WT=189, Lox=223) | |
| Aars2E16_geno_upper_Lox | |
| Aars2_geno_1_FOR | TGTTCATGGAGGAAGTGCCC |
| Aars2_geno_2_REV | CAGGAACCCCAACGGACAC |
| (3+4:WT=216, Lox=250) | |
| Aars2E16_geno_lower_Lox | |
| Aars2_geno_3_FOR | CAATGCACACCTCCGTGGAG |
| Aars2_geno_4_REV | TGCCGTTCCCCAATGATCAC |
| spanning whole (1+4:WT=438, 2lox=506) | |
| For validation PCR outside of the insertion region | |
| Aars2_geno_In14up_FOR_5 | GGAAGTGGCATGCTCCAAGA |
| Aars2_geno_In17dn_REV_6 | TCTCACACCTCACTGCAGTC |
| Aars2_geno_E12up_FOR_7 | GCTTCAGACCGAGGCTACCT |
| Aars2_geno_In19dn_REV_8 | GGCATGGCTGACTTCCTCTT |

**Supplemental Table 7. Antibodies and reagents.**

| Antibody | Host | Clone | Company | Cat# | Lot |
|---|---|---|---|---|---|
| **GAPDH** | Mouse | 1E6D9 | Proteintech | 60004-1 | 10013030 |
| **PCBP1** | Rabbit | EPR11049(B) | Abcam | ab168377 | GR121942-4 |
| **MF20** | Mouse | MF20 | DSHB | DSHB Hybridoma Product MF 20 | Supernatant produced in-house |
| **eIF2α** | Rabbit | Polyclonal | Cell Signaling | 9722 | 15 |
| **Phospho-eIF2α (Ser51)** | Rabbit | Polyclonal | Cell Signaling | 9721 | 21 |
| **AARS2** | Rabbit | Polyclonal | Abcam | ab197367 | GR206264-12 |
| **OxPhos Rodent WB Antibody Cocktail** | Mouse | Monoclonal | Abcam | STN-19467 | 2.1E+09 |
| **Endomucin** | Rat | | Santa Cruz | sc-65495 | B2720 |
| **Goat anti-Rabbit-AF594** | Goat | | Invitrogen | A11012 | 1892262, 1704538 |
| **Goat anti-Rabbit-AF488** | Goat | | Invitrogen | A11034 | 1937195 |
| **Goat anti-Rabbit-AF647** | Goat | | Invitrogen | A21245 | 2098544 |
| **Donkey anti-Mouse-AF488** | Donkey | | Invitrogen | A21202 | 2266877 |
| **Donkey anti-Mouse-AF594** | Donkey | | Invitrogen | A21203 | 2134005 |
| **Goat anti-Mouse-AF488** | Goat | | Invitrogen | A11029 | 1911843 |
| **Goat anti-Mouse-AF647** | Goat | | Invitrogen | A21235 | 43239A |
| **Donkey anti-Rabbit-AF594** | Donkey | | Invitrogen | A21207 | 2266563 |
| **Goat anti-Rat-AF647** | Goat | | Invitrogen | A21247 | 2251195 |

| | | | | | |
|---|---|---|---|---|---|
| **Donkey anti-Rat-AF488** | Donkey | | Invitrogen | A21208 | 538955 |
| **Hoechst-33342** | | | Invitrogen | 62249 | N/A |
| **DAPI (4',6-Diamidino-2-Phenylindole, Dihydrochloride)** | | | Invitrogen | D1306 | N/A |
| **Peroxidase AffiniPure Goat Anti-Rabbit IgG (H+L)** | | | Jackson Immunoresearch | 111-035-144 | |
| **Peroxidase AffiniPure Goat Anti-Mouse IgG (H+L)** | | | Jackson Immunoresearch | 115-035-146 | |

### RNAscope™ Probe

| Probe | Cat# |
|---|---|
| Mm-Hey2 | 404651 |
| Mm-Nppa | 418691 |
| Mm-Myl4 | 443801 |
| Mm-Myl7-C3 | 584271-C3 |
| Mm-Bmp10 | 415921 |
